# Independence of 3D chromatin conformation and gene regulation during *Drosophila* dorsoventral patterning

**DOI:** 10.1101/2020.07.07.186791

**Authors:** Elizabeth Ing-Simmons, Roshan Vaid, Mattias Mannervik, Juan M. Vaquerizas

## Abstract

The relationship between the 3D organisation of chromatin inside the nucleus and the regulation of gene expression remains unclear. While disruption of domains and domain boundaries can lead to mis-expression of developmental genes, acute depletion of key regulators of genome organisation, such as CTCF and cohesin, and major reorganisation of genomic regions have relatively small effects on gene expression. Therefore, it is unclear whether changes in gene expression and chromatin state drive chromatin reorganisation, or whether changes in chromatin organisation facilitate cell type-specific activation of genes and their regulatory elements. Here, using the *Drosophila melanogaster* dorsoventral patterning system as a model, we demonstrate the independence of 3D chromatin organisation and developmental gene regulation. We define tissue-specific enhancers and link them to expression patterns at the single-cell level using single cell RNA-seq. Surprisingly, despite tissue-specific differences in chromatin state and gene expression, 3D chromatin organisation is maintained across tissues. Our results provide strong evidence that tissue-specific chromatin conformation is not required for tissue-specific gene expression, but rather acts as an architectural framework to facilitate proper gene regulation during development.

---

Chromatin is highly organised within the nucleus into chromosome territories, compartments of active and inactive chromatin, self-interacting domains, and loops between specific loci. However, the relationship between the 3D organisation of chromatin and the regulation of gene expression remains unclear. There is much evidence that chromatin conformation is important for gene regulation: disruption of domains and domain boundaries can lead to mis-expression of developmental genes, contributing to developmental defects and cancer (Flavahan et al., 2016; Franke et al., 2016; Hnisz et al., 2016; Ibn-Salem et al., 2014; Lupiáñez et al., 2015; Spielmann et al., 2018). In addition, the general principles of 3D genome organisation are conserved across large evolutionary distances, as well as the chromatin conformation at specific loci (Fudenberg and Pollard, 2019; Harmston et al., 2017b; Krefting et al., 2018; Özdemir and Gambetta, 2019; Rowley et al., 2017; Vietri Rudan et al., 2015). Further evidence comes from the identification of interactions between promoters and their regulatory elements (Cruz-Molina et al., 2017; Ghavi-Helm et al., 2014; Hsieh et al., 2020; Jin et al., 2013; Krietenstein et al., 2020; Li et al., 2012; Sanyal et al., 2012; Weintraub et al., 2017), and the finding that forced enhancer-promoter looping is sufficient to activate transcription of some genes (Bartman et al., 2016; Deng et al., 2012; Deng et al., 2014; Morgan et al., 2017). However, in other cases direct enhancer-promoter contacts may be neither strictly required nor sufficient for gene activation (Alexander et al., 2019; Benabdallah et al., 2019; Chen et al., 2018; Heist et al., 2019). Furthermore, depletion of key regulators of genome organisation such as CTCF and cohesin has relatively small effects on gene expression (Nora et al., 2017; Rao et al., 2017; Schwarzer et al., 2017; Wutz et al., 2017), and genomic rearrangements are not always associated with changes in gene expression (Ghavi-Helm et al., 2019; Meadows et al., 2010; Williamson et al., 2019).

While multiple studies have shown differences in chromatin conformation between different cell types or tissues (Bonev et al., 2017; Chathoth and Zabet, 2019; Joshi et al., 2015; Kragesteen et al., 2018; Le Dily et al., 2014; Oudelaar et al., 2020; Schmitt et al., 2016), whether these changes are the cause or consequence of changes in gene expression is unclear. Therefore, a fundamental question arises as to whether changes in gene expression and chromatin state drive chromatin reorganisation, or whether changes in chromatin organisation facilitate cell type-specific activation of genes and their regulatory elements.

Embryonic development requires precise regulation of gene expression, making it an ideal context in which to investigate the relationship between gene regulation and chromatin organisation (Hug and Vaquerizas, 2018a). In particular, *Drosophila melanogaster* has long been used as a model organism for the study of development, and the key principles and factors involved in embryonic patterning are well understood (Wieschaus, 2016; Wolpert, 2016). Early *Drosophila* development involves a series of thirteen rapid, synchronous nuclear divisions, before the embryo becomes cellularised and undergoes zygotic genome activation (ZGA) at nuclear cycle (nc) 14 (Hamm and Harrison, 2018). We and others have previously shown that chromatin organisation in *Drosophila* is established at nc14, coincident with ZGA (Hug et al., 2017; Ogiyama et al., 2018). While a small number of genes are zygotically expressed prior to the major wave of ZGA (Lott et al., 2011), maternally provided cues are responsible for establishing the major anterior-posterior and dorsoventral axes (Ma et al., 2016; Stein and Stevens, 2014). Therefore, by ZGA, cells in different regions of the embryo contain different developmental transcription factors, have different patterns of chromatin accessibility (Cusanovich et al., 2018), and are primed to express different genes.

Cell fate along the dorsoventral (DV) axis is controlled by the nuclear concentration of the transcription factor Dorsal (Dl) (Hong et al., 2008; Stathopoulos and Levine, 2002), which peaks during nc14 (Reeves et al., 2012). Activation of the Toll signalling pathway on the ventral side of the embryo leads to high levels of Dl entering the nucleus, while Dl is excluded from the nucleus on the dorsal side (Stein and Stevens, 2014). Different levels of Dl concentration are responsible for the specification of different cell fates (Hong et al., 2008; Stathopoulos and Levine, 2002) (Fig. 1A). Maternal effect mutations in the Toll pathway lead to a uniform level of nuclear Dl across the embryo, making it possible to obtain females that produce a homogeneous population of embryos that consist entirely of presumptive mesoderm (*Toll^10B^*), neuroectoderm (*Toll^rm9/rm10^*), or dorsal ectoderm (*gd^7^*) (Fig. 1A). These embryos provide an excellent model system to study tissue-specific regulation during development, which has led to the discovery of key transcription factors, regulatory elements, and processes required for embryo patterning (Boija and Mannervik, 2016; Hong et al., 2008; Koenecke et al., 2016; Koenecke et al., 2017; Stathopoulos et al., 2002; Zeitlinger et al., 2007).

**Figure 1.**
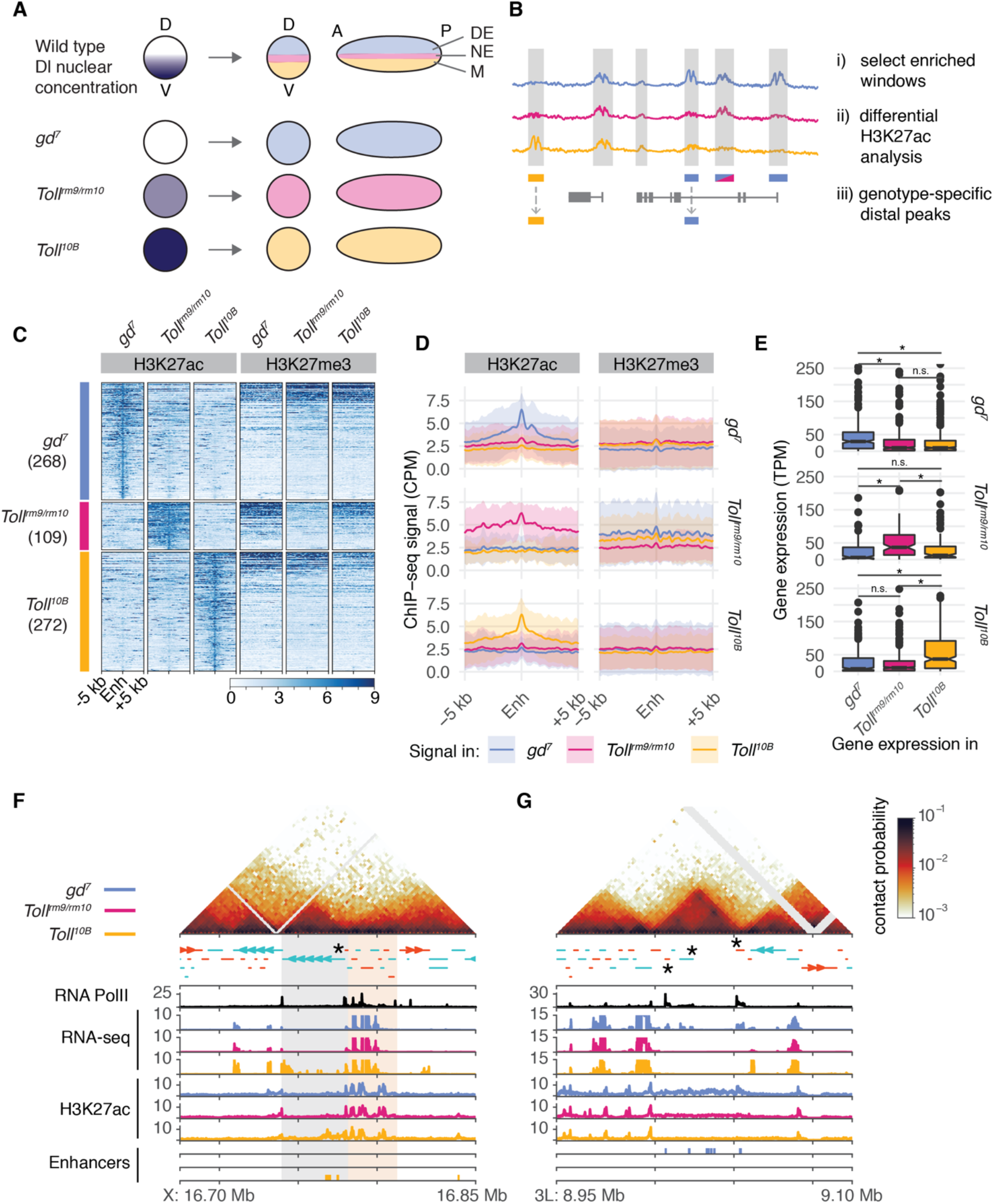
Identification of tissue-specific regulatory elements for dorsoventral patterning. **A.** Dorsoventral patterning of the Drosophila embryo is controlled by the nuclear concentration of Dl. High levels of nuclear Dl on the ventral side of the embryo produce mesoderm (ME, yellow), intermediate levels produce neuroectoderm (NE, pink), while nuclei without Dl will produce dorsal ectoderm (DE, blue). The *gd^7^*, *Toll^rm9/rm10^*, and *Toll^10B^* maternal effect mutations lead to embryos with uniform levels of nuclear Dl, which produce only dorsal ectoderm, neuroectoderm, and mesoderm, respectively. D, dorsal; V, ventral; A, anterior; P, posterior. **B.** Schematic representation of the identification of putative tissue-specific enhancers. In order to identify regions with tissue-specific H3K27ac, regions with enriched H3K27ac ChIP-seq signal compared to global background (grey shaded areas) were selected and pairwise comparisons between genotypes (*gd^7^*, blue; *Toll^rm9/rm10^*, pink; *Toll^10B^*, yellow) were performed using *csaw* (Lun and Smyth, 2014; Lun and Smyth, 2016). We selected regions that were enriched for H3K27ac in one genotype compared to both others, and removed regions that overlapped a promoter. **C**. Heatmaps of H3K27ac and H3K27me3 ChIP-seq signal at putative tissue-specific enhancers in dorsoventral mutant embryos. **D.** Average H3K27ac and H3K27me3 ChIP-seq signal at putative tissue-specific enhancers in dorsoventral mutant embryos. Shaded area represents +/− one standard deviation from the mean. **E.** Expression of genes associated with putative tissue-specific enhancers in dorsoventral mutant embryos. Top, genes associated with *gd^7^*-specific H3K27ac peaks; middle, genes associated with *Toll^rm9/rm10^*-specific H3K27ac peaks; bottom, genes associated with *Toll^10B^*-specific H3K27ac peaks. Asterisks indicate Wilcoxon rank-sum test p < 0.005. **F, G.** Examples of chromatin organisation at dorsoventral patterning genes (*if*, **F**; *Doc1*, *Doc2*, *Doc3*, **G**; these genes are labelled with asterisks). Top, normalised Hi-C contact probability maps, 2kb resolution, from 3-4 hpf control embryos (Hug et al., 2017). Positive strand genes are shown in orange and negative strand genes are shown in blue. RNA PolII-Ser5p ChIP-seq from late nc14 control embryos is shown in black (Blythe and Wieschaus, 2015). RNA-seq and H3K27ac ChIP-seq from 2-4 hpf dorsoventral mutant embryos is shown in blue (*gd^7^*), pink (*Toll^rm9/rm10^*), and yellow (*Toll^10B^*) ((Koenecke et al., 2016), this study). Tissue-specific putative enhancers identified as described above are shown as colour-coded bars. The grey shaded area in **F** represents a domain containing a developmentally regulated gene; the orange shaded area highlights a region containing housekeeping genes.

In this study, we use *Drosophila* DV patterning as a model system to investigate the relationship between tissue-specific gene regulation and 3D chromatin organisation. We focus on the cellular blastoderm stage, approximately 2-3 hours post fertilisation (hpf), which is coincident with nc14, establishment of chromatin organisation, and the onset of ZGA. We identify putative regulatory elements involved in DV patterning, and show that their target genes are developmentally regulated and have a distinct chromatin organisation compared to housekeeping genes. We find that while there are clear differences in chromatin state and overall gene expression between *gd^7^*, *Toll^rm9/rm10^*, and *Toll^10B^* embryos, there is still significant heterogeneity in gene expression at the single-cell level. However, these tissue-specific differences in chromatin state and gene expression are not associated with tissue-specific 3D chromatin organisation. Together, these results provide strong evidence that tissue-specific chromatin conformation is not required for tissue-specific gene expression. Rather, our findings indicate that the organisation of the genome into 3D chromatin domains acts as an architectural framework to facilitate correct regulation of gene expression during development.

## RESULTS

### Identification of regulatory elements and genes involved in dorsoventral patterning

In order to identify candidate tissue-specific regulatory elements involved in dorsoventral patterning, we carried out genome-wide differential peak identification for the active chromatin mark H3K27ac, using ChIP-seq data from 2-4 hpf embryos, including ChIP-seq data for *Toll^rm9/rm10^* embryos generated in this study and for *gd^7^* and *Toll^10B^* embryos from (Koenecke et al., 2016) (Fig. 1B). This identified 268 regions enriched for H3K27ac in *gd^7^* compared to both *Toll^rm9/rm10^* and *Toll^10B^*, 109 regions specifically enriched in *Toll^rm9/rm10^*, and 272 regions specifically enriched in *Toll^10B^* (Fig. 1C). These putative tissue-specific enhancers are depleted for the repressive chromatin mark H3K27me3 in the tissue in which they are enriched for H3K27ac (Fig. 1C, 1D), providing further evidence for their tissue specificity. By requiring significant enrichment in one genotype compared to both other genotypes, we select a highly stringent set of regions with tissue-specific increases in H3K27ac. These putative enhancers overlap with genomic regions that have been shown to drive expression in the expected regions of the embryo (Fig. S1D, E). Next, we assigned putative enhancers to target genes using a combination of gene expression data, linear genomic proximity, and chromatin conformation data (see Methods), and verified that genes assigned to tissue-specific candidate enhancers have significantly higher expression in the tissue where the enhancer is active (Fig. 1E, all p < 0.001). We conclude that the identified regions represent a stringent set of candidate enhancers associated with regulation of dorsoventral patterning.

### Developmentally regulated genes have a distinct regulatory landscape compared to housekeeping genes

We next assessed the chromatin conformation landscape around these tissue-specific regulatory elements and their target genes. Using Hi-C data from 3-4 hpf Drosophila embryos (Hug et al., 2017), we observed that dorsoventral patterning genes and their regulatory elements are located within the same self-interacting domains (Fig. 1F, grey shaded region; Fig. 1G). These domains are larger than domains not associated with developmentally regulated genes (Fig. S1A, B) (mean size 98 kb compared to 67 kb, p = 3.47 x 10^−14^). This is in contrast to housekeeping genes, which are enriched at the boundaries between domains and in small domains (Hug et al., 2017; Sexton et al., 2012; Ulianov et al., 2016) (Fig. 1F, orange shaded region). In addition, domains containing developmentally regulated genes are significantly more likely to overlap with the large regions of high non-coding sequence conservation known as Genomic Regulatory Blocks (Engström et al., 2007; Harmston et al., 2017b; Kikuta et al., 2007) (Fig. S1C). These results are robust to different definitions of tissue-specific enhancers (Fig. S2) and emphasize the distinct organisation of developmentally regulated and housekeeping genes in the *Drosophila* genome.

### Single-cell expression analysis reveals heterogeneity in gene expression within dorsoventral mutant embryos

While the *gd^7^*, *Toll^rm9/rm10^*, and *Toll^10B^* maternal effect mutants have long been used as models to analyse tissue-specific regulation during dorsoventral patterning (Hong et al., 2008; Stathopoulos et al., 2002; Zeitlinger et al., 2007), the extent of cell fate conversion at the single-cell level within these embryos is unknown. Anterior-posterior patterning mechanisms are still active and RNA *in situ* hybridisation experiments suggest that cell fate conversion may be incomplete (Stathopoulos and Levine, 2002). In order to assess the heterogeneity of gene expression and cell identities within these embryos, we carried out single-cell gene expression analysis using the 10X Genomics Chromium platform. We used embryos at 2.5-3.5 hpf to target the late cellular blastoderm stage when dorsoventral patterning has been established (Reeves et al., 2012; Sandler and Stathopoulos, 2016). Clustering of single-cell expression profiles from wild type and mutant embryos showed good concordance with bulk RNA-seq (Fig. S3A) and identified 15 clusters representing different cell identities within the embryo (Fig. 2A, Fig. S3C). Two of these represent cells from embryos in the earlier stages of cellularisation, due to the timed collection. Other clusters represent mesoderm, ectoderm, amnioserosa, and terminal regions of the embryo, in addition to more specific cell populations such as pole cells, haemocytes, and trachea precursors. A full list of clusters and cluster marker genes is available in Table S2. Visualisation of single-cell gene expression profiles from *gd^7^*, *Toll^rm9/rm10^*, and *Toll^10B^* embryos revealed that specific clusters are depleted in these mutant embryos (Fig. 2B, Fig. S3B-C). Cells from clusters representing mesoderm cell fates are almost completely absent in *gd^7^* and *Toll^rm9/rm10^* embryos, and subsets of ectoderm cells are missing in each of the mutants (Fig. 2B, Fig. S3B-C).

**Figure 2.**
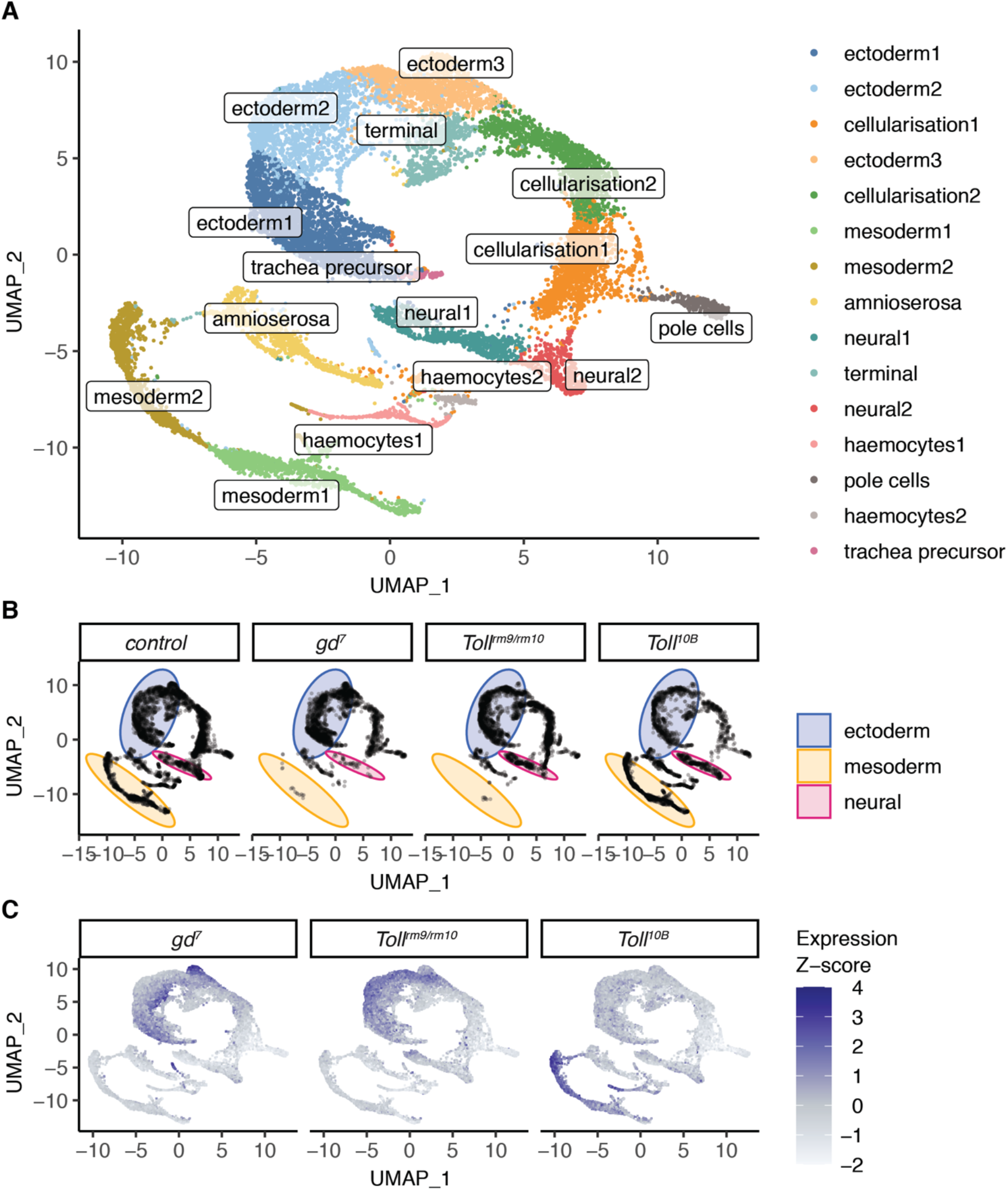
Single-cell RNA-seq analysis of gene expression during dorsoventral patterning. **A.** Clustering of single-cell gene expression profiles from 2.5-3.5 hpf embryos reveals clusters corresponding to distinct cell populations. **B.** Single cell gene-expression profiles separated by cell origin (control, *gd^7^, Toll^rm9/rm10^*, or *Toll^10B^* embryos). Certain clusters are depleted in the dorsoventral mutant embryos. Blue, area of graph corresponding to ectoderm clusters; pink, neural; yellow, mesoderm. **C.** Single-cell expression of genes associated with putative tissue-specific enhancers. Colour is mapped to the Z-score of average expression of genes assigned to the groups of tissue-specific enhancers identified in Figure 1.

In order to further dissect the ectoderm clusters and identify cells corresponding to dorsal ectoderm and neuroectoderm, we visualised expression of genes assigned to tissue-specific enhancers (Fig. 2C) and known dorsal ectoderm, neuroectoderm, and mesoderm marker genes (Fig. S3C, (Karaiskos et al., 2017)). This revealed that the ectoderm clusters contain distinct subpopulations of cells expressing dorsal ectoderm markers and neuroectoderm markers. These subpopulations correspond to the regions of the cell distribution which are depleted in *Toll^rm9/rm10^* and *gd^7^* respectively (Fig. 2B, compare distributions in the ‘ectoderm’ region). Therefore, while the mutant embryos evidently still consist of a mixture of cell identities, certain cell fates are lost. Overall, these results highlight that these embryos have a significant level of cell-to-cell heterogeneity. Importantly, the loss of specific cell fates combined with the tissue-specific enhancer usage shown above supports the use of these embryos to model dorsoventral patterning perturbations.

### Major features of chromatin organisation are conserved across tissues

We next asked how differential usage of regulatory elements and differential gene expression relates to chromatin conformation during dorsoventral patterning. To do so, we generated Hi-C datasets for *gd^7^*, *Toll^rm9/rm10^*, *Toll^10B^,* and control embryos at the cellular blastoderm stage, at 2kb resolution (Table S3).

Systematic comparison of the Hi-C datasets across genotypes revealed that on average, chromatin conformation is similar across datasets (Fig. 3). Saddle plots reveal similar strength of compartmentalisation in control and *gd^7^*, *Toll^rm9/rm10^*, and *Toll^10B^* embryos (Fig. 3A, B). We next analysed overall self-interacting domain strength using domains identified in control 3-4 hpf embryos as a reference (Fig. 3C). While domain strength is weaker in cellular blastoderm embryos than at 3-4 hpf, the strength is similar across all genotypes, suggesting that the vast majority of domains and domain boundaries are present in all tissues. We obtained similar conclusions when we examined chromatin loop strengths, using loops from Kc167 cells (Cubeñas-Potts et al., 2017) as a reference (Fig. 3D, E), indicating that loops are conserved across tissues. Finally, we analysed genome-wide contact probability decay with distance (*P(s)*). A shallow slope at distances < 100 kb reflects local chromatin compaction into domains, while the flattening of the slope around separation distances of 1 Mb indicates compartment formation ((Gassler et al., 2017; Naumova et al., 2013; Schwarzer et al., 2017), Fig. 3F). We also examined the derivative of *P(s)*, since this can highlight differences in the strength of domain formation (Fig. 3G) (Gassler et al., 2017). These analyses reveal differences in these profiles at distances > 5 Mb, which correspond to genomic rearrangements on balancer chromosomes present in a subset of *Toll^rm9/rm10^* and *Toll^10B^* embryos (Fig. S4). Combined, our results demonstrate that overall genome organisation at the level of compartments, domains, and chromatin loops is similar across tissues in cellular blastoderm embryos.

**Figure 3.**
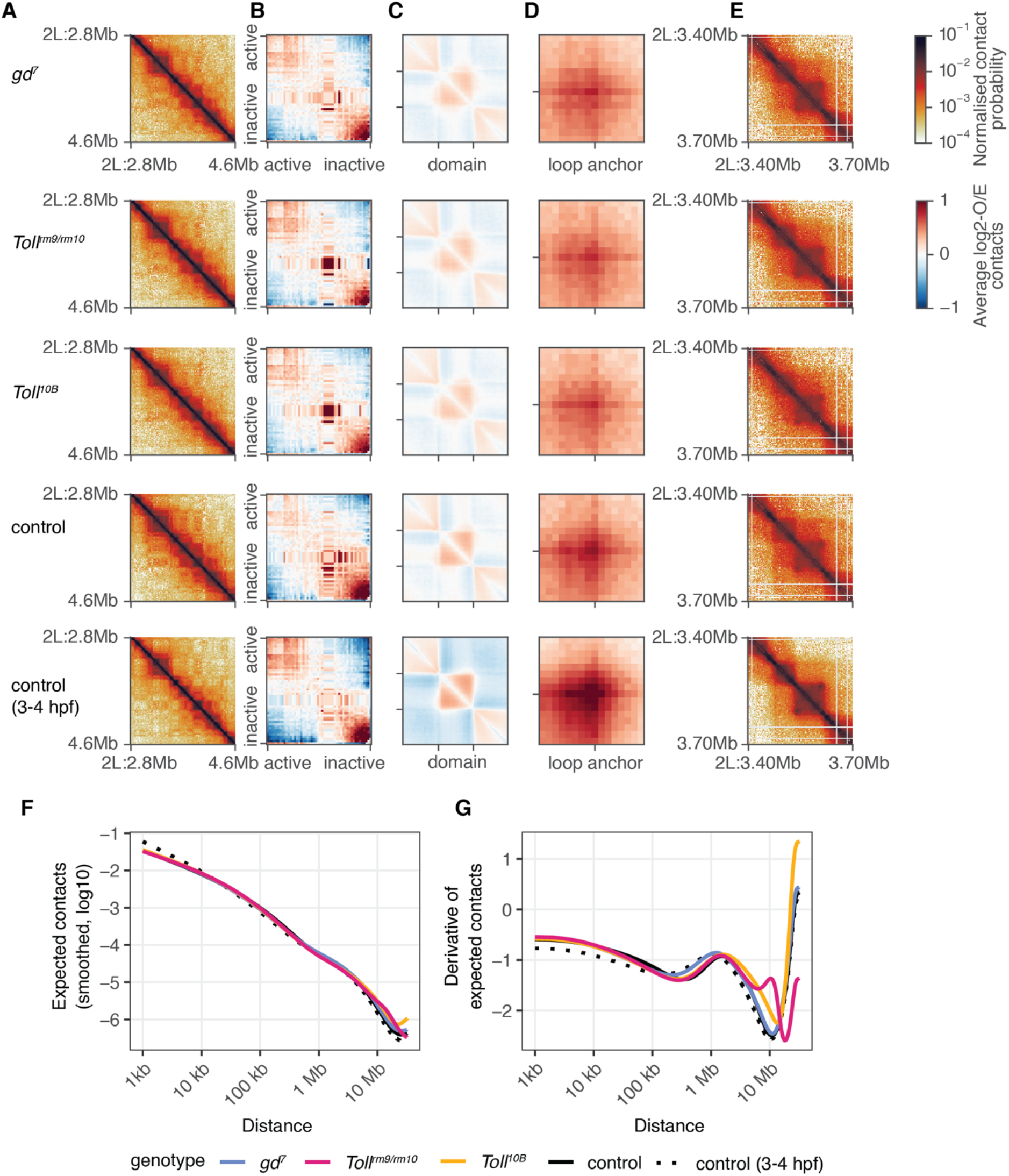
Global chromatin conformation along the dorsoventral axis. **A.** Chromatin conformation for a 1.8 Mb region of chromosome 2L in *gd^7^*, *Toll^rm9/rm10^*, *Toll^10B^*, and control embryos at the cellular blastoderm stage, and for control embryos at 3-4 hpf using data from (Hug et al., 2017). **B.**‘Saddle-plot’ representing genome-wide average chromatin compartmentalisation. Active regions tend to interact with other active regions (top left), while inactive regions interact with other inactive regions (bottom right). **C.** Aggregate analysis of domains identified using Hi-C data from 3-4 hpf embryos at 2kb resolution. **D.** Aggregate analysis of chromatin loops identified in (Cubeñas-Potts et al., 2017). **E.** Chromatin conformation for a 300 kb region of chromosome 2L, showing domains and a loop. **F.** Average contact probability at different distances for control (black), *gd^7^* (blue), *Toll^rm9/rm10^* (pink), and *Toll^10B^* (yellow) embryos. **G.** The derivative of the expected contact probability by distance, highlighting differences between samples at far-cis distances due to the presence of balancer chromosomes in the *Toll^rm9/rm10^* and *Toll^10B^* embryos.

### Chromatin conformation at developmentally regulated genes is similar across tissues despite differences in gene expression and chromatin state

In order to systematically assess chromatin conformation across the genome and identify regions with differences, we used CHESS (Galan, Machnik et al., in revision), a novel approach for differential chromatin conformation detection based on computer vision techniques. Briefly, Hi-C submatrices are compared genome-wide between pairs of datasets to produce a similarity score and a signal to noise ratio for each pair of genomic windows (see Methods). Using this approach, we compared control and mutant embryos at the cellular blastoderm stage at 5kb resolution and with a 500 kb window size. As a reference, we compared control cellular blastoderm stage data from this study with Hi-C data from nc14 embryos from (Hug et al., 2017). We subtracted this reference score from the score for each control-mutant comparison in order to identify regions with differences in genome organisation between control and mutant embryos. This analysis revealed that most regions across the genome do not display significant differences in 3D chromatin organisation between *gd^7^*, *Toll^rm9/rm10^*, and *Toll^10B^* embryos (Fig. 4A, Fig. S4, Fig. S5). This agreed with visual examinations of control-mutant difference matrices (Fig. 4E). The subset of regions that do display strong changes in chromatin organisation between genotypes can be attributed to genomic rearrangements present on balancer chromosomes in a subset of the *Toll^rm9/rm10^* and *Toll^10B^* embryos, rather than correlating with the locations of genes that are differentially expressed (Fig. 4A-B, Fig. S4–S6). To further investigate the relationship between changes in chromatin conformation and gene expression, we analysed CHESS similarity scores in windows containing genes that are differentially expressed in pairwise comparisons between the mutant embryos (Koenecke et al., 2016) compared to other genomic windows (Fig. 4C). This revealed a lack of association between differential gene expression and differential chromatin structure genome wide. To further validate these observations, we visually assessed genome organisation in *gd^7^*, *Toll^rm9/rm10^*, and *Toll^10B^* embryos at known differentially expressed dorsoventral patterning genes (Fig. 4E, Fig. S6). Examination of chromatin conformation and chromatin data at these regions does not indicate any differences in domain organisation, boundary formation, or loop formation. For example, the *Doc1*, *Doc2* and *Doc3* genes, which are highly expressed in *gd^7^* and required for amnioserosa differentiation and dorsolateral ectoderm patterning (Reim, 2003), lie in a well-insulated domain that contains multiple *gd^7^*-specific putative enhancers and is enriched for H3K27me3 in all three mutants at 2-4 hpf, although to a lesser extent in *gd^7^* (Fig. 4E). There is no evidence of changes in the insulation of this domain in *gd^7^*, where the genes are active, compared to the other datasets, nor is there any change in its internal structure such as changes in interactions between enhancers and the target gene promoters. Similar conclusions were obtained from examining additional loci, including the *pnr* locus (expressed in *gd^7^*) and the *NetA*/*NetB*, *if*, and *sna* loci which are active in *Toll^10B^* (Fig. S6). Together, these results suggest that tissue-specific gene expression and enhancer activity do not necessarily involve changes in domain organisation.

**Figure 4.**
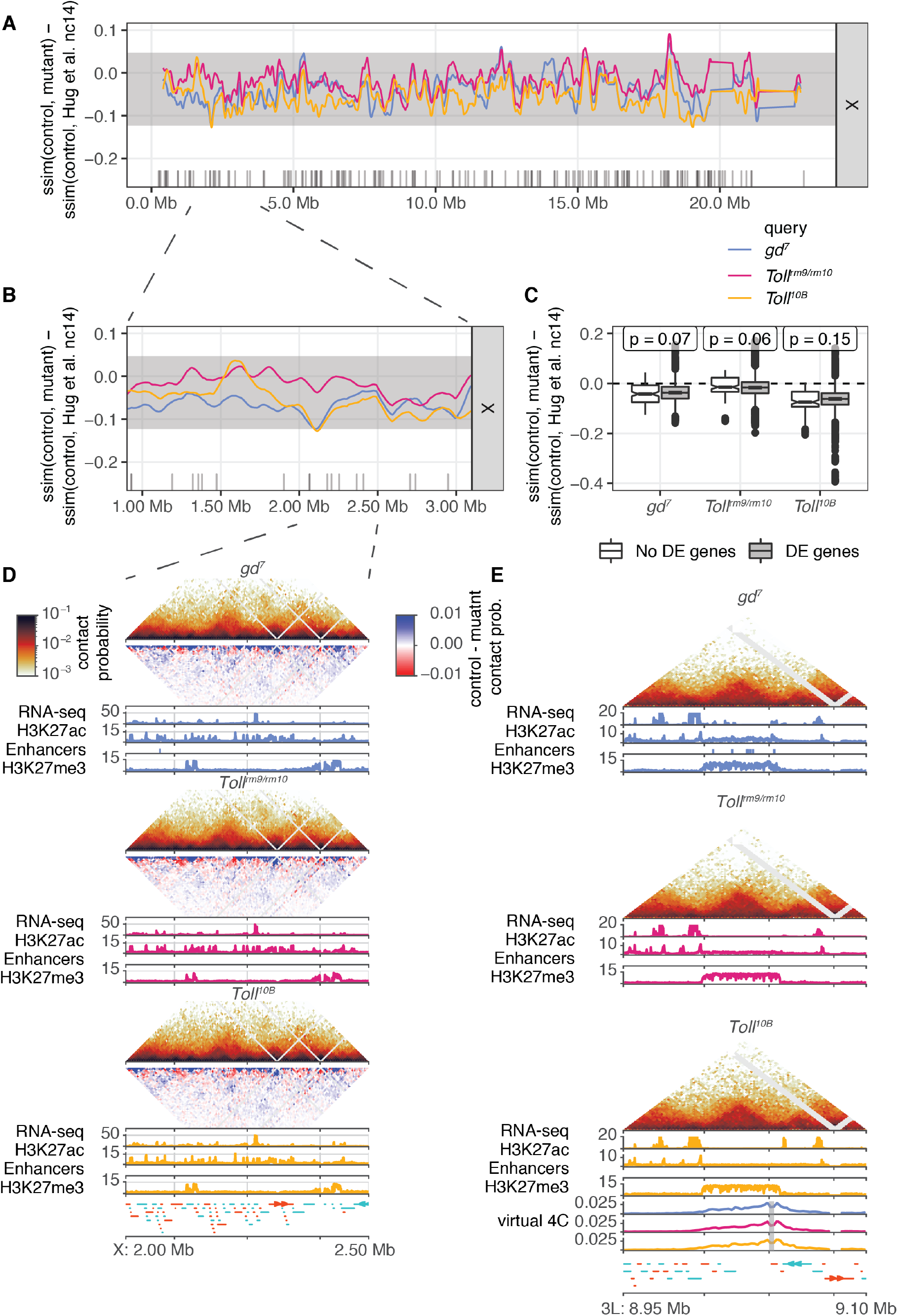
Chromatin conformation is not affected by tissue-specific gene expression. **A.** CHESS (Galan, Machnik et al., in revision) similarity scores were calculated between mutant and control embryo Hi-C datasets, using 5kb resolution and a 500 kb window size. As a reference, similarity scores were calculated between control embryo Hi-C data and the nc14 Hi-C data from (Hug et al., 2017). The difference between this reference similarity score and the similarity scores between mutant and control embryos for chromosome X is shown (blue, *gd^7^*; pink, *Toll^rm9/rm10^*; yellow, *Toll^10B^*). Similarity score differences around zero represent regions where chromatin conformation is similar between control and mutant, while negative values represent regions where there are greater differences between control and mutant than between control and the Hug et al. data. Shaded area represents similarity score differences within two standard deviations of the genome-wide mean. Grey ticks represent the positions of genes that are differentially expressed between dorsoventral mutant embryos (Koenecke et al., 2016). Other chromosomes are shown in Figure S4. **B.** Example of differences in CHESS similarity scores for a 2 Mb region on chromosome X. Differences in similarity score do not strongly correlate with the positions of differentially expressed genes. Hi-C data for a subset of this region is shown in **D**. **C.** Boxplots of differences in similarity score for genomic windows with or without genes that are differentially expressed (DE) in pairwise comparisons between dorsoventral mutant embryos (Koenecke et al., 2016). There are no significant differences between windows with and without DE genes (Wilcoxon rank sum test, all p > 0.05). **D.** Tissue-specific chromatin data for a subset of the region on chromosome X shown in **C** above. For each genotype, top, normalised Hi-C contact probability maps and Hi-C difference maps at 5kb resolution. Hi-C difference maps are calculated as *contact probability in control – contact probability in mutant*; red indicates regions with increased contact probability in embryos of the mutant genotype and blue indicates decreased contact probability. Bottom, RNA-seq (Koenecke et al., 2016), H3K27ac and H3K27me3 ChIP-seq data ((Koenecke et al., 2016), this study). Tissue-specific putative enhancers identified as described above are shown as colour-coded bars beneath the corresponding H3K27ac ChIP-seq track. Lower panel, gene annotations. Positive-strand genes are shown in orange and negative strand genes are shown in blue. See also additional example loci in Figure S5. **E.** Tissue-specific chromatin data for the region around the *Doc1*, *Doc2* and *Doc3* genes shown in Figure 1E. For each genotype, top, normalised Hi-C contact probability maps, at 2kb resolution; middle, RNA-seq data (Koenecke et al., 2016); bottom, H3K27ac and H3K27me3 ChIP-seq data ((Koenecke et al., 2016), this study). Tissue-specific putative enhancers identified as described above are shown as colour-coded bars beneath the corresponding H3K27ac ChIP-seq track. Bottom, “virtual 4C” tracks for each genotype representing interactions of a 2kb region around the promoter of *Doc1*; gene annotations. Positive-strand genes are shown in orange and negative strand genes are shown in blue. See also additional example loci in Figure S6.

Finally, traditional models of gene regulation by enhancers predict that interactions between regulatory elements and their target promoters increase upon tissue-specific gene expression (Alberts et al., 2014). While such changes were not apparent upon visual examination (Fig. 4D, Fig. S6), we performed aggregate analysis to systematically determine whether subtle changes in interaction strength manifest at these loci. This revealed that there is no significant increase in interaction frequency between enhancers and their target promoters in the tissue in which the enhancer is active (Fig. S7).

Taken together, our results provide strong evidence for the independence of tissue-specific gene expression and chromatin conformation during dorsoventral patterning.

## DISCUSSION

Our results indicate that despite significant differences in chromatin state and gene expression between tissues, cell type-specific gene regulation does not require cell type-specific chromatin conformation. Nevertheless, developmentally regulated genes and enhancers are organised into chromatin domains. We suggest that this organisation plays a permissive role to facilitate the precise regulation of developmental genes.

Maternal effect mutations in the Toll signalling pathway lead to embryos that lack the usual patterning of the DV axis (Stein and Stevens, 2014). These embryos have long been used as a system to study the specification of mesoderm (*Toll^10B^*), neuroectoderm (*Toll^rm9/rm10^*), and dorsal ectoderm (*gd^7^*) cell fates and the regulation of tissue-specific gene expression (Boija and Mannervik, 2016; Koenecke et al., 2016; Koenecke et al., 2017; Stathopoulos et al., 2002; Zeitlinger et al., 2007). However, these embryos are still under the influence of anterior-posterior patterning signals and do not show completely uniform cell identities (Stathopoulos et al., 2002). We sought to investigate heterogeneity of cell identity at the single-cell level by using single-cell gene expression profiling. This revealed that certain cell types are indeed maintained in all three Toll pathway mutants, including pole cells and other terminal region cell identities, haemocytes, and trachea precursor cells (Fig. 2). However, heterogeneity of gene expression is reduced in the mutants, as shown by the loss of cells assigned to mesoderm clusters in *gd^7^* and *Toll^rm9/rm10^* embryos, and the depletion of ectoderm subsets in each of the mutants. These datasets showcase the advantages of measuring cellular heterogeneity at the single cell level and will provide a useful resource for further characterisation of these embryos and investigation of the regulation of DV patterning.

Although the *gd^7^*, *Toll^rm9/rm10^*, and *Toll^10B^* embryos still have heterogeneous gene expression profiles, nevertheless there are clear differences in chromatin state and overall gene expression between them (Boija and Mannervik, 2016; Koenecke et al., 2016; Stathopoulos et al., 2002; Zeitlinger et al., 2007). We expanded on previous studies by identifying putative enhancers specific to neuroectoderm in addition to dorsal ectoderm and mesoderm. This allowed the identification of tissue-specific putative enhancer-gene pairs, which correspond well with known dorsoventral patterning enhancers and genes that are differentially expressed across the dorsoventral axis. These regulatory elements and their target genes are located inside chromatin domains, distinct from the enrichment of housekeeping genes at domain boundaries (Cubeñas-Potts et al., 2017; Hou et al., 2012; Hug et al., 2017; Ramírez et al., 2018; Sexton et al., 2012; Ulianov et al., 2016). This is in line with previous results that suggest that 3D chromatin domains act as regulatory domains (Despang et al., 2019; Harmston et al., 2017b; Ibrahim and Mundlos, 2020; Le Dily and Beato, 2015; Symmons et al., 2014).

Previous studies have produced conflicting results about whether tissue-specific enhancer-promoter interactions are correlated with tissue-specific activation of gene expression. Studies using 3C approaches have found evidence of enriched enhancer-promoter interactions (Hsieh et al., 2020; Jin et al., 2013; Krietenstein et al., 2020; Li et al., 2012; Sanyal et al., 2012), which may precede (Cruz-Molina et al., 2017; Ghavi-Helm et al., 2014) or correlate with (Bonev et al., 2017; Oudelaar et al., 2020) transcriptional activation. However, several recent studies in mammals as well as *Drosophila* provide evidence that stable enhancer-promoter contacts are not required for gene activation (Alexander et al., 2019; Benabdallah et al., 2019; Chen et al., 2018; Heist et al., 2019; Mir et al., 2018). Our results indicate stable chromatin conformation across tissues during *Drosophila* ZGA, and we found no evidence for widespread enrichment of interactions between enhancers and their target promoters. This is in line with “transcriptional hub” models, in which transient or indirect contacts with a regulatory element are sufficient to activate transcription (Furlong and Levine, 2018; Lim et al., 2018; Mir et al., 2018; Tsai et al., 2017; Yokoshi and Fukaya, 2019), such as through the formation of phase-separated condensates (Cho et al., 2018; Chong et al., 2018; Sabari et al., 2018). While a subset of long-range enhancer-promoter pairs do form stable interactions that are enriched above local background (Ghavi-Helm et al., 2014), these are not likely to be the primary mechanism of promoter regulation during *Drosophila* development. Many stable loops in the *Drosophila* genome are instead associated with Polycomb repression (Eagen et al., 2017; Ogiyama et al., 2018).

Previous studies have also not provided conclusive evidence whether changes in domain and compartment organisation are a cause or consequence of changes in chromatin and transcriptional activity. Some studies suggested that domains are widely conserved across different tissues and even different species (Dixon et al., 2012; Schmitt et al., 2016; Vietri Rudan et al., 2015). However, there are also numerous examples of changes in domain structures across cell types and differentiation time points (Bonev et al., 2017; Joshi et al., 2015; Kragesteen et al., 2018; Kruse et al., 2019; Le Dily et al., 2014). However, it is important to note that detection of domains and boundaries in different samples may be confounded by sequencing depth, analysis resolution, noise, and the presence of so-called “sub-domain” structure. In *Drosophila*, Hi-C maps from anterior and posterior embryo halves showed no differences (Stadler et al., 2017), and it has been proposed that active chromatin, especially at broadly expressed genes, is responsible for partitioning the genome into domains (Rowley et al., 2017; Ulianov et al., 2016). Rowley et al. (Rowley et al., 2017) proposed that compartmentalisation of active and inactive chromatin, at the level of individual genes, underlies the formation of insulated chromatin domains. This model predicts that when a developmentally regulated gene is active, its domain would merge with, or have increased interactions with, neighbouring domains containing active genes, for example broadly expressed housekeeping genes. In contrast to these predictions, we find that domain organisation is conserved across tissues, even in cases where there are significant changes in the local chromatin state and gene expression (Fig. 3 and Fig. S4). This suggests that, similarly to mammalian domain architecture, additional factors such as insulator proteins modulate domain organisation in *Drosophila* (Mateo et al., 2019; Moretti et al., 2020).

Together, our results indicate that differential chromatin organisation is not a necessary feature of cell-type specific gene expression. We propose that chromatin organisation into domains instead provides a scaffold or framework for the regulation of developmental genes during and after the activation of zygotic gene expression (Fig. 5, left and middle). This increases the frequency of contacts between regulatory elements and their target genes, as in models where enhancers scan the domain for their target promoter, and allows for cell type-specific activation of gene expression (Fig. 5, right). Feedback effects such as downstream modification of chromatin state and additional mechanisms including looping between Polycomb-bound elements and segregation of active and inactive chromatin then act as layers on top of the initially established domain structure.

**Figure 5.**
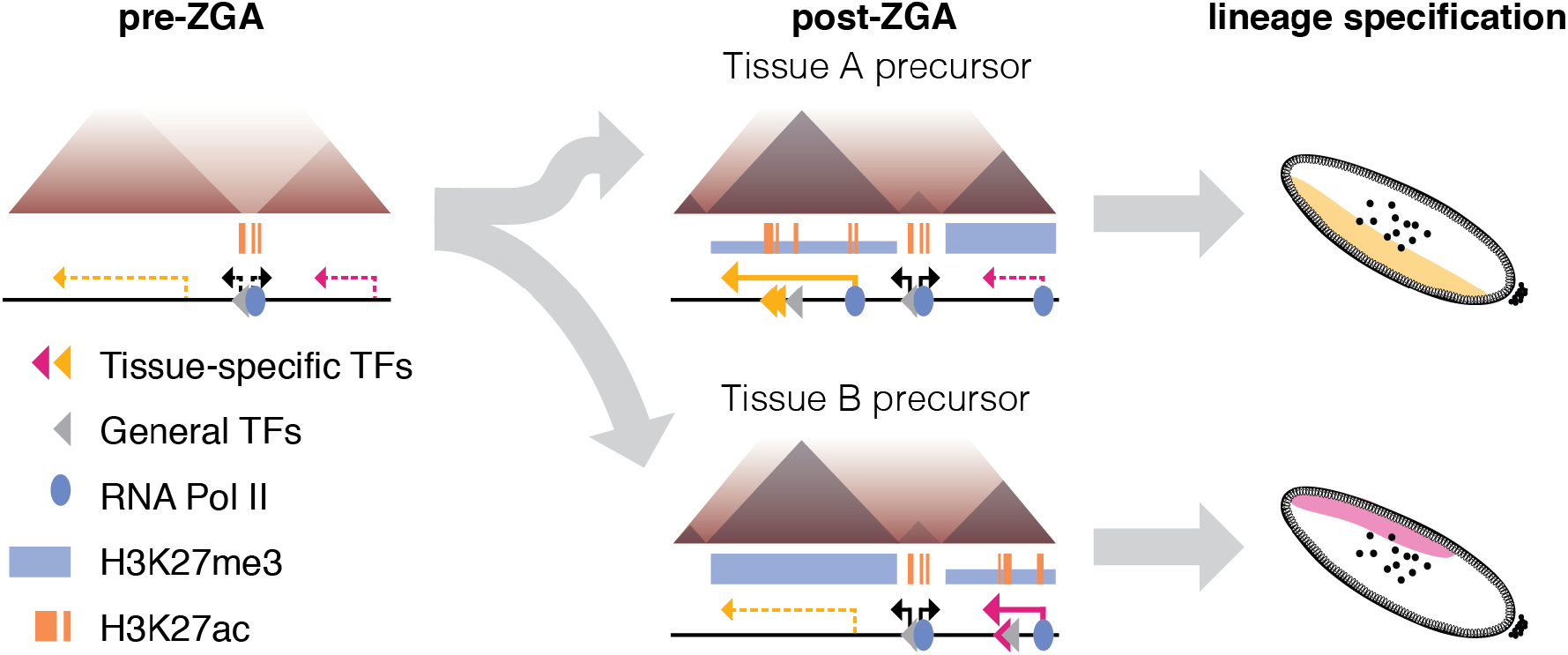
Model of the relationship between chromatin conformation and developmentally regulated gene expression. Before ZGA, the genome is unstructured (left), with domain boundaries appearing at a subset of regions associated with binding of RNA PolII and Zelda. Chromatin domains are established at ZGA, and domain structure is the same across tissues with different gene expression and transcription factor binding (middle). Differential activity of regulatory elements in the context of the same chromatin conformation leads to different patterns of gene expression in the developing embryo (right). Thick and thin blue bars represent high and low levels of H3K27me3 respectively; dashed lines represent inactive genes while solid lines represent actively transcribed genes.

## Author Contributions

Conceptualisation: E.I.-S. and J.M.V.; Formal analysis: E.I.-S.; Funding acquisition: E.I.-S., M.M., and J.M.V.; Investigation: E.I.-S. and R.V.; Resources: R.V. and M.M.; Supervision: M.M. and J.M.V.; Visualisation: E.I.-S.; Writing – original draft: E.I.-S.; Writing – review and editing: E.I.-S., R.V., M.M., and J.M.V.

## Acknowledgements

We thank the core facility at Novum, BEA – Bioinformatics and Expression Analysis, which is supported by the board of research at the Karolinska Institute and the research committee at the Karolinska hospital for help with sequencing. We are grateful to Christine Rushlow for critical reading of the manuscript, and members of the Vaquerizas lab for helpful discussions and feedback.

## METHODS

### Drosophila stock maintenance and embryo collection

yw; eGFP-PCNA flies used as controls for Hi-C and the first scRNA-seq control experiment were kindly provided by S.A. Blythe and E. Wieschaus (Blythe and Wieschaus, 2016) and maintained on standard cornmeal-agar food. The *w^1118^* flies used for the second scRNA-seq control experiment and the Toll mutant fly stocks *gd^7^/winscy hs-hid, Toll^10B^*/*TM3 e Sb Ser/OR60* and *Toll*^rm9/rm10^/*TM6 e Tb Sb* were grown on potato mash-agar food. All fly stocks were incubated at 25 °C with a 12-hour light/dark cycle.

The embryos representing presumptive dorsal ectoderm were collected from *gd^7^* homozygous flies. One day old larvae laid by *gd^7^/winscy hs-hid* were heat shocked for 1.5 hr at 37 °C twice with 24hr interval to eliminate gd^*7*^ heterozygous animals. Embryos from *Toll^10B^*/*TM3 e Sb Ser* or *Toll^10B^/OR60* heterozygous females represented presumptive mesoderm. *Toll*^rm9^*/Toll*^rm10^ trans-heterozygous females were used for collecting presumptive neuroectoderm embryos.

We adapted the fixation and sorting procedure described in (Blythe and Wieschaus, 2015) for *in situ* Hi-C (Hug and Vaquerizas, 2018b; Rao et al., 2014). Following a pre-collection period of at least one hour, fly embryos were collected on yeasted 0.4 % acetic acid agar plates or apple juice plates at 25 °C. After 1 hour of collection, the embryos on the plate were incubated at 25 °C for 2 hours. Embryos were dechorionated for 2 min in 2.6 % sodium hypochlorite, rinsed with water, and transferred to vials containing 2 mL of PBS, 0.5 % Triton X-100 and 6 mL of heptane. Cross-linking was initiated by adding 100 µl of 37 % formaldehyde, followed by vigorous shaking. After 10 min, samples were spun at 500 g for 1 min and the upper heptane layer was removed. 15 min after the start of fixation, 5 mL of PBS, 0.5 % Triton X-100, 125 mM glycine was added to the embryos, followed by vigorous shaking for 1 min. The embryos were rinsed 3 times with PBS, 0.5 % Triton X-100. Embryos were sorted in small batches under a light microscope, based on morphology, to select embryos of the appropriate developmental stage and remove damaged embryos or embryos with abnormal morphology. Sorted embryos were aliquoted so that a single tube contained enough embryos for one experiment, then flash-frozen in liquid nitrogen and stored at −80 °C. We used 30-60 embryos for each *in situ* Hi-C experiment.

For single-cell RNA-seq, we adapted the collection and methanol fixation procedures described in (Alles et al., 2017; Karaiskos et al., 2017). Following a pre-collection period of at least one hour, fly embryos were collected on yeasted apple juice plates at 25 °C. After 1 hour of collection, the embryos on the plate were incubated at 25 °C for 2.25 hours. Embryos were dechorionated for 2 min in 2.6 % sodium hypochlorite, rinsed with water, and suspended in PBS, 0.5 % Triton X-100. Embryos were rinsed with cell culture grade DPBS without Ca^2+^ and Mg^2+^ to remove residual detergent, and placed on ice at precisely 3.5 hours after the start of collection. Embryos were resuspended in 500 µl ice cold dissociation buffer (cell culture grade DPBS without Ca^2+^ and Mg^2+^, 0.04 % BSA) and dissociated with a clean metal pestle. Cells and tissue fragments were pelleted at 500 x g for 5 min at 4 °C, then gently resuspended in 100 µl Trypsin-EDTA 0.25 % and incubated for 3 minutes. After 3 minutes Trypsin was quenched by adding 1 mL cell culture grade DPBS without Ca^2+^ and Mg^2+^, 10 % FCS. Cells were pelleted at 1000 g for 5 min at 4 °C, then resuspended in 500 µl dissociation buffer, pelleted again and resuspended in 100 µl dissociation buffer. A 10 µl aliquot of cells was kept and counted using an improved Neubauer chamber or a Luna2 cell counter. To fix cells, 4 volumes of 100 % methanol, pre-chilled at −20 °C, were slowly added to the cells. Fixed cells were stored at −80 °C and used within 3 days.

### ChIP-seq

2-4 hr old *Toll*^rm9^*/Toll*^rm10^ embryos for ChIP sequencing were collected and fixed as described above for Hi-C. Fixed embryos were flash-frozen in liquid nitrogen and stored at −80 °C until further use. Frozen embryos were homogenized in sonication buffer (50 mM HEPES, 140 mM NaCl, 1 mM EDTA, 1 % Triton, 0.1 % sodium deoxycholate, 0.1 % SDS and protease inhibitor) using a Dounce homogenizer. The sample were spun at 4000 g for 5 min and the pellet containing the intact nuclei was resuspended in same buffer supplemented with 0.5 % *N*-Lauroylsarcosine and SDS to final concentration of 0.5 %. The chromatin was sheared to fragment size in the range of 200–500 bp using a Bioruptor (Diagenode). The solubilized chromatin fraction was cleared by centrifugation and used for immunoprecipitation after diluting 5 times with sonication buffer. Immunoprecipitation with either 2 µg of H3K27ac (Abcam, ab4729) or 5 µg of H3K27me3 (Abcam, ab6002) antibody was carried out on chromatin corresponding to 20-25 µl of embryos at 4 °C overnight. Chromatin-antibody complexes were captured for at least 3 hr using a mix of Protein A and G Dynabeads (Invitrogen). The captured immunoprecipitated complex were washed 10 min each with sonication buffer (50 mM HEPES, 140 mM NaCl, 1 mM EDTA, 1 % Triton, 0.1 % sodium deoxycholate, 0.1 % SDS), WashA (as sonication buffer, but with 500 mM NaCl), WashB (20 mM Tris pH 8, 1 mM EDTA, 250 mM LiCl, 0.5 % NP-40, 0.5 % sodium deoxycholate) and TE. After the washes Dynabeads with bound chromatin-antibody complexes were resuspended in 100 µl TE supplemented with 20 mg/ml RNase A and incubated at 50 °C for 30 min. Cross-linking was reversed by adding Tris pH 8.0 and SDS to a final concentration of 50 mM and 0.1% respectively and heating at 68 °C for at least 4 hr. Protein digestion was carried out by Proteinase K treatment at 55 °C for 2 hr, followed by purifying ChIP DNA using ChIP DNA Clean & Concentrator kit (Zymo research #D5205). ChIP-seq libraries were prepared on the ChIP DNA eluted in 60 µl of DNA elution buffer, using the NEBNext Ultra II DNA Library Prep Kit (NEB). ChIP samples were single-end (1 × 75 bp) sequenced on Illumina NextSeq platform at BEA core facility, Stockholm.

### scRNA-seq

We performed scRNA-seq using the 10X Genomics Chromium Single Cell 3’ Reagents v3, according to the manufacturer’s instructions (Rev B). Methanol-fixed cells were spun at 3000 g at 4 °C for 5 min and resuspended in 500 µl DPBS + 0.04 % BSA to rehydrate. Rehydrated cells were counted using a Luna2 cell counter, and the volume used for library preparation was chosen for a targeted recovery of 5000 cells. Libraries were sequenced on an Illumina NextSeq 500, using paired-end sequencing with read 1 length 28 cycles, index read length 8 cycles, and read 2 length 91 cycles.

### Hi-C

We performed *in situ* Hi-C according to the protocol in (Hug and Vaquerizas, 2018b; Hug et al., 2017), using MboI as the restriction enzyme, with minor modifications for optimisation for low input according to (Díaz et al., 2018).

### ChIP-seq analysis

ChIP-seq reads were mapped to the dm6 genome using Bowtie2 (version 2.3.3.1 (Langmead and Salzberg, 2012)). Mapped reads were filtered to remove alignments with quality scores less than 30, as well as secondary and supplementary alignments. PCR duplicates were marked using sambamba (version 0.6.8, (Tarasov et al., 2015)). Coverage tracks were generated using the bamCoverage tool from deepTools (version 3.2.0, (Ramírez et al., 2014)) with the following parameters: “-of bigwig --binSize 10 --normalizeUsing CPM --extendReads 200 --ignoreDuplicates --minMappingQuality 30” and keeping only reads from chromosomes X, 2L, 2R, 3L, 4, and Y. ChIP-seq peaks were called using MACS2 (version 2.2.6, (Feng et al., 2012)) with the following parameters: “--nomodel --extsize 147 -g dm” or “--nomodel --extsize 147 -g dm --broad --min-length 500 --max-gap 200” for broad peaks. We used merged input samples for each genotype as the controls for all peak calling, due to a lack of sample-matching information for the published datasets that were re-analysed.

### Bulk RNA-seq analysis

RNA-seq reads were quantified using Salmon (1.1.0, (Patro et al., 2017)) and the Flybase r6.30 transcripts. Salmon was used in mapping-based mode, with the following parameters: "-l A --validateMappings –seqBias". For visualisation purposes, RNA-seq reads were also aligned to the dm6 genome, using Hisat2 (version 2.1.0, (Kim et al., 2019)). Mapped reads were filtered to remove alignments with quality scores less than 30, as well as secondary and supplementary alignments. PCR duplicates were marked using sambamba (version 0.6.8, (Tarasov et al., 2015)). Coverage tracks were generated using the bamCoverage tool from deepTools (version 3.2.0, (Ramírez et al., 2014)) with the following parameters: “-of bigwig --binSize 10 --normalizeUsing CPM --extendReads 200 --ignoreDuplicates --minMappingQuality 30” and keeping only reads from chromosomes X, 2L, 2R, 3L, 4, and Y.

We used tximport (version 1.14.2, (Soneson et al., 2016)) to import quantifications from Salmon into R (3.6.3) and estimate transcripts per million values. We carried out pairwise differential expression analysis between *gd^7^*, *Toll^rm9/rm10^*, and *Toll^10B^* using DESeq2 (version 1.26.0, (Love et al., 2014)) with default parameters.

### Identification of candidate tissue-specific enhancers

In order to identify tissue-specific enhancers, we first carried out pairwise differential H3K27ac signal analysis using csaw (version 1.20.0, (Lun and Smyth, 2014; Lun and Smyth, 2016)) and edgeR (version 3.28.1, (Robinson et al., 2010)). We used 2000 bp windows for the background calculations and selected 150 bp windows with a 3-fold enrichment over the background. Windows were merged using the parameters “tol = 100” and “max.width = 5000”. Merged regions with an FDR < 0.05 and with a consistent direction of change across all windows were selected for downstream analysis. Candidate tissue-specific enhancers were defined by taking the intersection of regions identified as enriched for H3K27ac in each genotype compared to both others.

We validated the putative enhancers by comparing to enhancers identified in previous studies. 7 of 22 dorsal ectoderm enhancers identified from a literature search (Koenecke et al., 2016) overlap with our *gd^7^*-specific enhancers, while 13 of 37 mesoderm enhancers overlap our *Toll^10B^*-specific enhancers (Fig. S1C). The relatively low overlap can be explained by the fact that many literature enhancers have H3K27ac signal in *Toll^rm9/rm10^* as well as either *gd^7^* or *Toll^10B^*. Putative enhancers were also overlapped with regions tested for enhancer activity in *Drosophila* embryos by (Kvon et al., 2014). Regions (“tiles”) tested by Kvon et al. that were active in at least one tissue and timepoint were lifted over to dm6 from dm3. 114 putative enhancers overlapped a total of 124 tiles by at least 1bp. Out of 27 *gd^7^* enhancers which overlap tiles that are active in stage 4-6 or stage 7-8, 21 are active in either dorsal ectoderm or amnioserosa precursors/subsets. Out of 16 *Toll^rm9/rm10^* enhancers that overlap tiles that are active in stage 4-6 or stage 7-8, 14 are active in brain or ventral nerve cord precursors, procephalic ectoderm, or ventral ectoderm. Out of the 30 *Toll^10B^* enhancers that overlap tiles that are active in stage 4-6 or stage 7-8, 28 are active in mesoderm precursors/subsets.

Enhancer heatmaps were made using the plotHeatmap tool from deepTools (version 3.2.0, (Ramírez et al., 2014)). Overlaps between different enhancer sets were visualised using UpSetR (version 1.4.0, (Conway et al., 2017; Lex et al., 2014)).

### Assignment of candidate enhancers to target genes

We defined “housekeeping genes” as genes that have at least ‘low’ expression in all stages and tissues according to Flybase RNA-seq data (1867 genes). We filtered the set of genes from the Flybase 6.30 transcripts to remove these housekeeping genes, as well as any genes with an average TPM < 1 in the *gd^7^*, *Toll^rm9/rm10^*, and *Toll^10B^* bulk RNA-seq. Candidate tissue-specific enhancers were assigned to target genes using the following rules: first, we assigned any enhancers that overlapped a single transcript to that gene. Next, we assigned enhancers to the closest promoter that was not separated from the enhancer by a domain boundary (using consensus boundaries from 3-4 hpf embryos, see below). The remaining enhancers were assigned to the closest promoter within the same domain, or, if they were not inside a domain, to the closest promoter.

### scRNA-seq analysis

We used CellRanger (version 3.1.0) to produce fastq files for the scRNA-seq data and to align, filter, and quantify reads based on the BDGP6.22 genome release (Ensembl 98) to produce feature-barcode matrices. We imported the filtered matrices into R using DropletUtils (version 1.6.1, (Griffiths et al., 2018)), and performed additional quality control analysis using scater (version 1.14.6, (McCarthy et al., 2017)). Doublets were identified using scDblFinder (version 1.1.8, (Germain et al., 2020)), with an estimated doublet rate of 3.9%, and removed. Normalisation for library size across cells was performed using scater and scran (version 1.14.6, (Lun et al., 2016)) using the “deconvolution” approach described in (Amezquita et al., 2019), in which cells are pre-clustered and size factors estimated using the calculateSumFactors() function.

Downstream analysis was carried out using Seurat (version 3.1.4, (Butler et al., 2018; Stuart et al., 2019)). The VST method was used to select the top 3000 variable features for each sample, then all datasets were integrated using the control dataset with the highest number of cells (replicate 1) as the reference dataset, and the first 30 dimensions. We performed clustering using the Shared Nearest Neighbour approach implemented in the Seurat functions FindNeighbors and FindClusters, using the first 12 dimensions from PCA, a k.param value of 60, and a clustering resolution of 0.5. These parameters were chosen because they produced clusters that were stable to small variations in the parameter values.

We carried out differential expression analysis using the Seurat function FindMarkers in order to identify genes with higher expression in a cluster compared to all other cells, and in pairwise comparisons. We carried out Gene Ontology enrichment analysis on the results marker gene sets using the enrichGO function from clusterProfiler (version 3.14.3, (Yu et al., 2012)), and simplified the results to remove semantically similar terms using the simplify function from clusterProfiler with the Wang method and a similarity threshold of 0.7. These marker gene sets and enriched GO terms, along with expression of known markers for embryonic cell populations, were used to assign putative cluster identities.

In order to quantify the average expression of particular gene sets in Figure 2C and Figure S3C, we calculated the sum of expression of those genes per cell, and then expressed this as a Z-score across all cells.

Pooled scRNA-seq reads from all barcodes were analysed using Salmon as described above.

### Hi-C analysis

Hi-C data was analysed using FAN-C (version 0.8.28) (Kruse et al., 2020). Paired-end reads were scanned to identify ligation junctions, split at ligation junctions if any were present, and then aligned independently to the dm6 genome using BWA-MEM (version 0.7.17-r1188) (Li and Durbin, 2009). Aligned reads were filtered to retain only uniquely-aligned reads with a mapping quality of at least 3. Reads were then paired based on read names and assigned to restriction fragments. “Inward” and “outward” reads separated by less than 1 kb, representing likely unligated fragments and self-ligated fragments respectively, were removed. In addition, we removed PCR duplicates, reads mapping more than 500 bp from a restriction site, and self-ligations where both reads map to the same fragment.

We generated two biological replicate datasets for each genotype, which showed high similarity. Therefore, we pooled biological replicates to reach 2 kb resolution. Matrices created from merged biological replicates were binned and filtered using FAN-C default parameters to remove bins with coverage less than 10 % of the median coverage. Normalisation was performed using Knight-Ruiz matrix balancing (Knight and Ruiz, 2013). Expected contacts are calculated as the average contacts at each genomic distance separation. Hi-C data was visualised using plotting tools from FAN-C and using HiGlass (Kerpedjiev et al., 2018).

### Domain and boundary identification

The insulation score was calculated as described in (Crane et al., 2015), using FAN-C, for 2 kb and 5 kb resolution matrices, each with window sizes of 4, 6, 8, and 10 bins. Domain boundaries were calculated from the insulation score with a delta parameter of 3 and filtered to keep only boundaries with a boundary score of at least 0.7. Consensus boundaries for each sample were created by overlapping boundaries called at the two different resolutions and four different window sizes, and keeping those boundaries which were identified using at least four out of the total eight parameter combinations. Domains were created by pairing boundaries, and domains less than 10 kb or more than 500 kb in size were removed.

### Hi-C aggregate analysis

Aggregate compartment, domain, and loop plots were created using FAN-C. Compartments were identified using the first eigenvector of the correlation matrix of the normalised Hi-C data, using GC content to orient the eigenvector. The compartment eigenvector for the 3-4 hpf Hi-C data from Hug et al. 2017 was used as the reference for the aggregate compartment plots (“saddle plots”) (Flyamer et al., 2017; Gassler et al., 2017; Imakaev et al., 2012; Kruse et al., 2020). Domain aggregates were also created using the domains identified in the Hug et al. 3-4 hpf Hi-C data. Loop aggregates were created using the loops identified in Kc167 cells by (Cubeñas-Potts et al., 2017). Similar results were obtained using loops from (Eagen et al., 2017; Stadler et al., 2017) (not shown).

We constructed a BEDPE file of putative enhancer-promoter interactions, considering all unique transcript start sites for assigned target genes. Interactions with a separation of at least 10 kb were used to create enhancer-promoter aggregate plots.

### Hi-C similarity score analysis with CHESS

We used CHESS (Comparison of Hi-C Experiments using Structural Similarity) (Galan, Machnik et al., in revision) to compare Hi-C data from embryos of different genotypes. Briefly, CHESS treats Hi-C interaction matrices as images and applies the concept of the structural similarity index (SSIM), which is widely used in image analysis. We applied CHESS to 5 kb resolution Hi-C matrices using windows of 500 kb and a step size of 5 kb to produce similarity scores for pairwise Hi-C comparisons. Hi-C data from stage 5 control embryos was used as the reference dataset, and compared to data from stage 5 dorsoventral mutant embryos and nc14 control embryos from Hug et al. 2017. In order to identify regions of the genome with significant changes between the reference and query datasets, we selected regions with a SSIM Z-score less than −2, and a signal-to-noise ratio Z-score of at least 1.

### Statistics and visualisation

Statistical tests were carried out in R (version 3.6.3) and visualisation was performed using the ggplot2 package (Wickham, 2016). Box plots are defined with boxes spanning the first to third quartiles. The whiskers extend from the box to the smallest/largest values no further than 1.5 times the interquartile range (IQR) away from the box. The notches extend 1.58 * IQR / sqrt(n) from the median.

### Data availability

The Hi-C, scRNA-seq, and ChIP-seq data produced in this study have been submitted to ArrayExpress.

## SUPPLEMENTARY FIGURES

**Figure S1.**
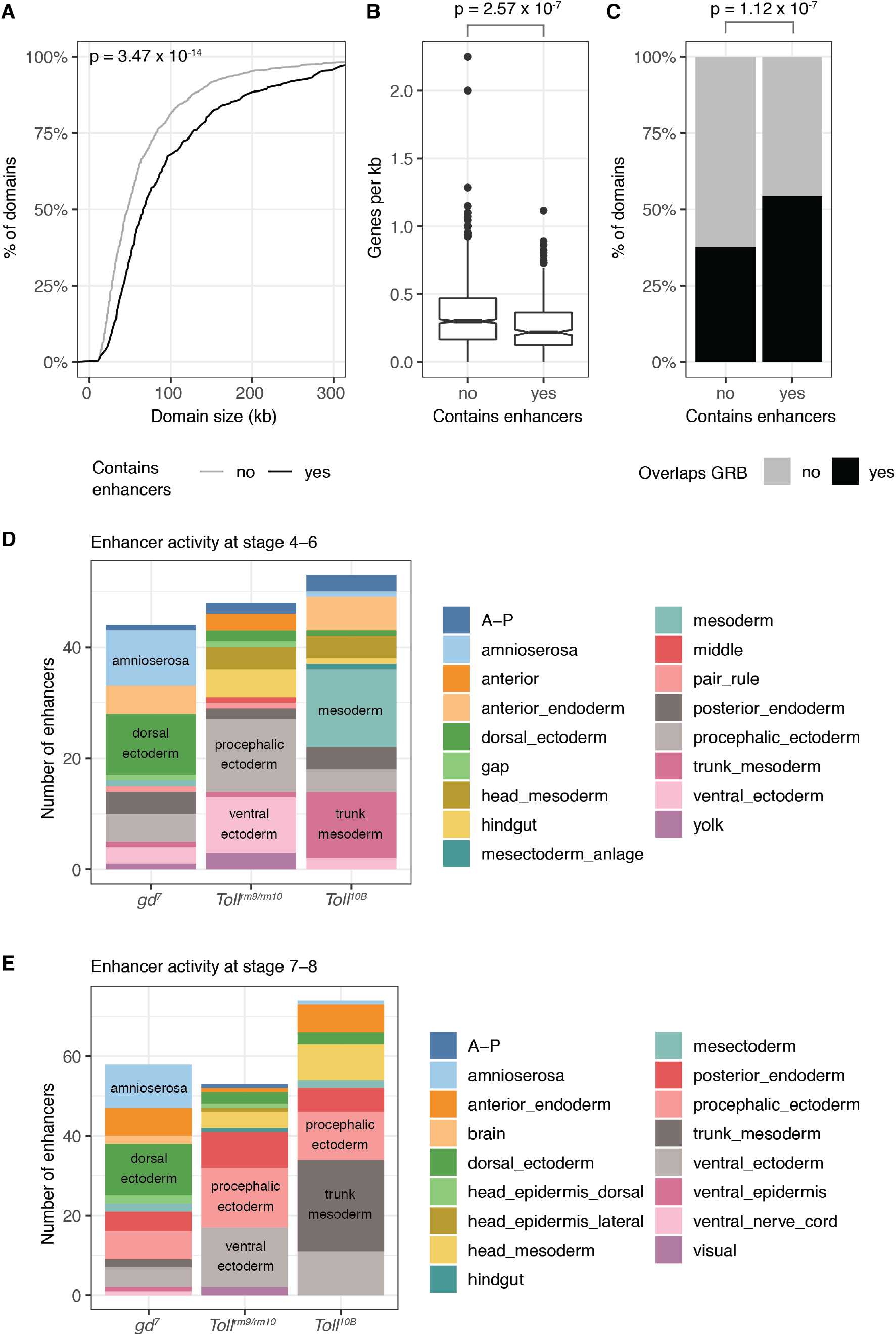
Properties of domains containing tissue-specific putative enhancers and validation of tissue-specific enhancer activity. **A.** Size of domains containing putative tissue-specific enhancers (black) and those without tissue-specific enhancers (grey). Wilcoxon rank sum test p = 3.47 x 10^−14^. **B.** Genes per kilobase inside domains containing putative tissue-specific enhancers and those without tissue-specific enhancers. Wilcoxon rank sum test p = 2.57 x 10^−7^. **C.** Overlap of chromatin domains with Genomic Regulatory Blocks (GRBs) from (Harmston et al., 2017a). Chi-squared test p = 1.12 x 10^−7^. **D, E.** Putative enhancers active in *gd^7^*, *Toll^rm9/rm10^*, and *Toll^10B^* embryos overlap with enhancers tested by (Kvon et al., 2014) and are enriched for enhancers that drive expression in relevant regions of the embryo at stages 4-6 (**D**) and 7-8 (**E**). The two largest categories are labelled for each stacked bar.

**Figure S2.**
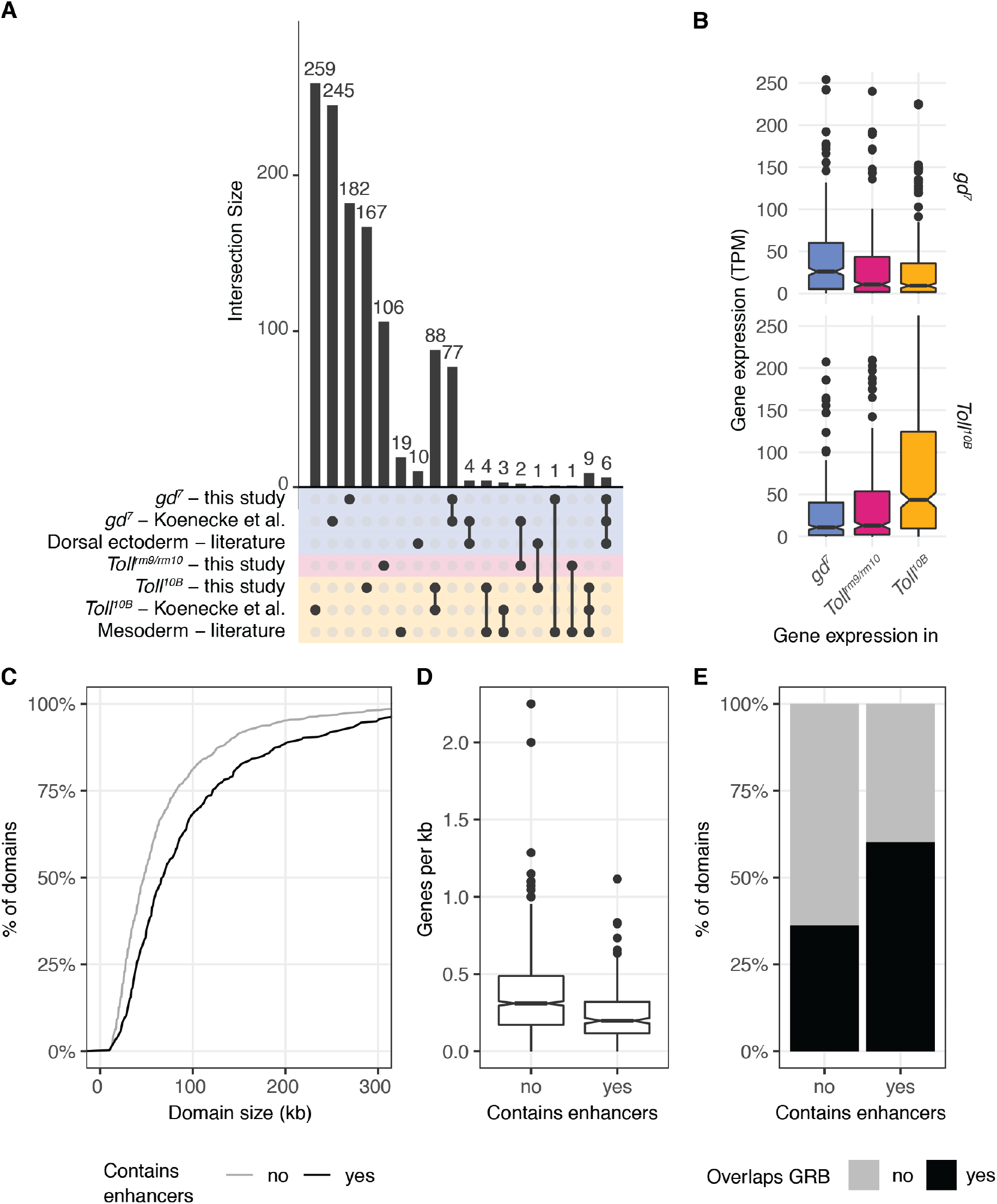
Properties of domains containing tissue-specific putative enhancers are robust to different enhancer definitions. **A.** UpSet plot showing the overlap between the putative tissue specific enhancers identified in this study, those identified in (Koenecke et al., 2016), and mesoderm and dorsal ectoderm enhancers identified from a literature search by (Koenecke et al., 2016). The lower panel represents the groups of genomic regions considered (y axis), and intersections between these groups are shown by joined dots. The bars represent the sizes of these intersections. For example, there are 6 putative dorsal ectoderm enhancers that are identified in this study, identified by Koenecke et al., and previously identified from a literature search (rightmost column). **B.** Expression of genes associated with putative enhancers identified by Koenecke et al. in *gd^7^* and *Toll^10B^* mutant embryos. Top, genes associated with *gd^7^*-specific putative enhancers; bottom, genes associated with *Toll^10B^*-specific putative enhancers. Asterisks indicate Wilcoxon rank sum test p < 0.001. **C.** Size of domains containing putative tissue-specific enhancers identified by (Koenecke et al., 2016) (black) and those without tissue-specific enhancers (grey). Wilcoxon rank sum test p = 7.12 x 10^−13^. **D.** Genes per kilobase inside domains containing putative tissue-specific enhancers identified by (Koenecke et al., 2016) and those without tissue-specific enhancers. Wilcoxon rank sum test p = 7.67 x 10^−16^. **E.** Overlap of chromatin domains with Genomic Regulatory Blocks (GRBs) from (Harmston et al., 2017a). Chi-squared test p = 3.14 x 10^−14^.

**Figure S3.**
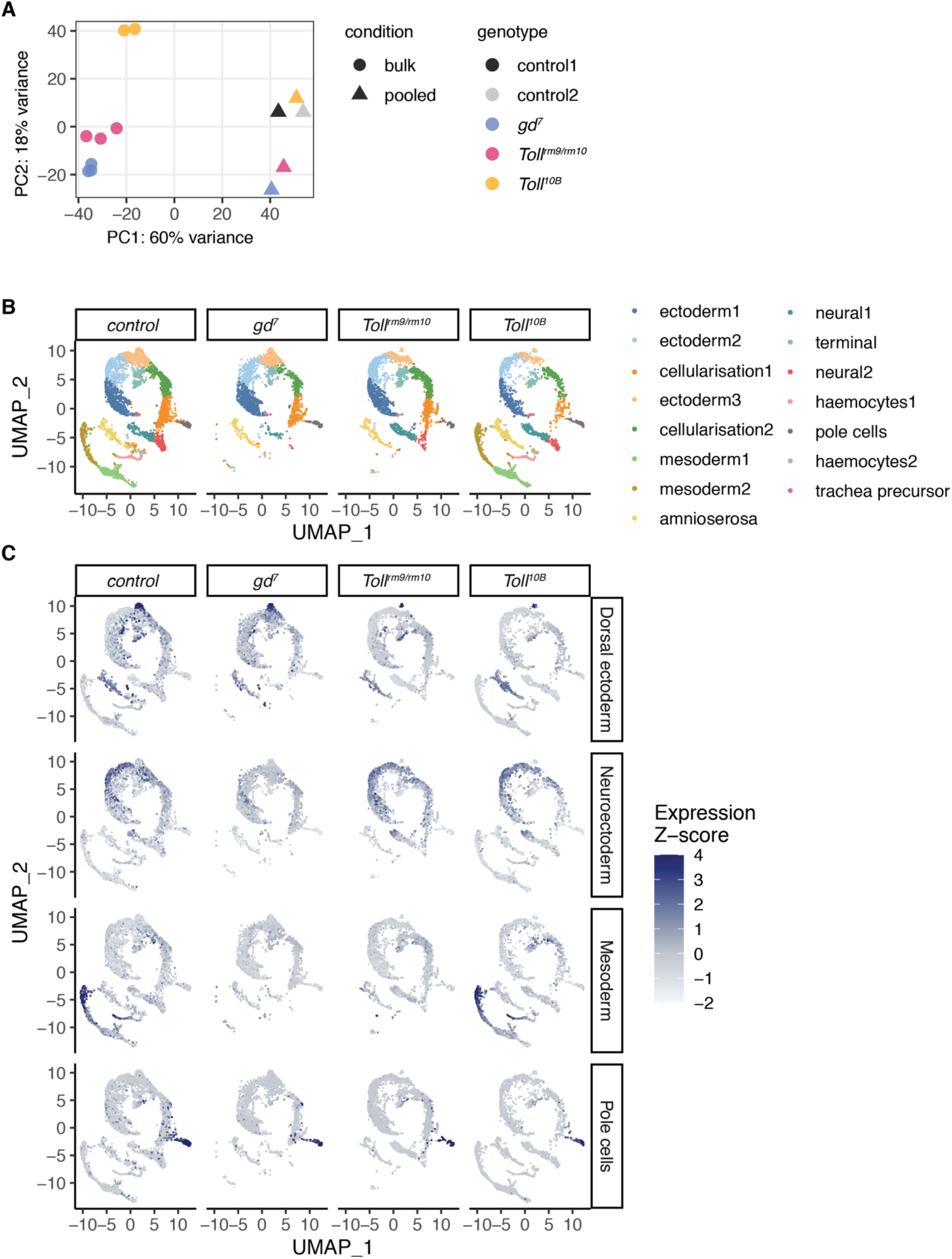
Validation of scRNA-seq data. **A.** PCA of pooled single-cell RNA-seq data and bulk RNA-seq data. The first principal component separates the methods while the second principal component separates the genotypes. Replicate control single-cell RNA-seq experiments cluster together, demonstrating robustness. **B.** Clustering of single-cell gene expression profiles from 2.5-3.5 hpf embryos reveals clusters corresponding to distinct cell populations, as in Fig 2A but separated by genotype of origin. **C.** Expression of tissue-specific marker genes across single cells from different genotype origins. Marker genes for dorsal ectoderm (*Ance, CG2162, Doc1, Doc2, egr, peb, tok, ush, zen*), neuroectoderm (*ac, brk, CG8312, l(1)sc, mfas, Ptp4E, sog, SoxN, vnd*), mesoderm (*CG9005, Cyp310a1, GEFmeso, ltl, Mdr49, Mes2, NetA, ry, sna, stumps, twi, wgn, zfh1*), and pole cells (*pgc*) were obtained from (Karaiskos et al., 2017).

**Figure S4.**
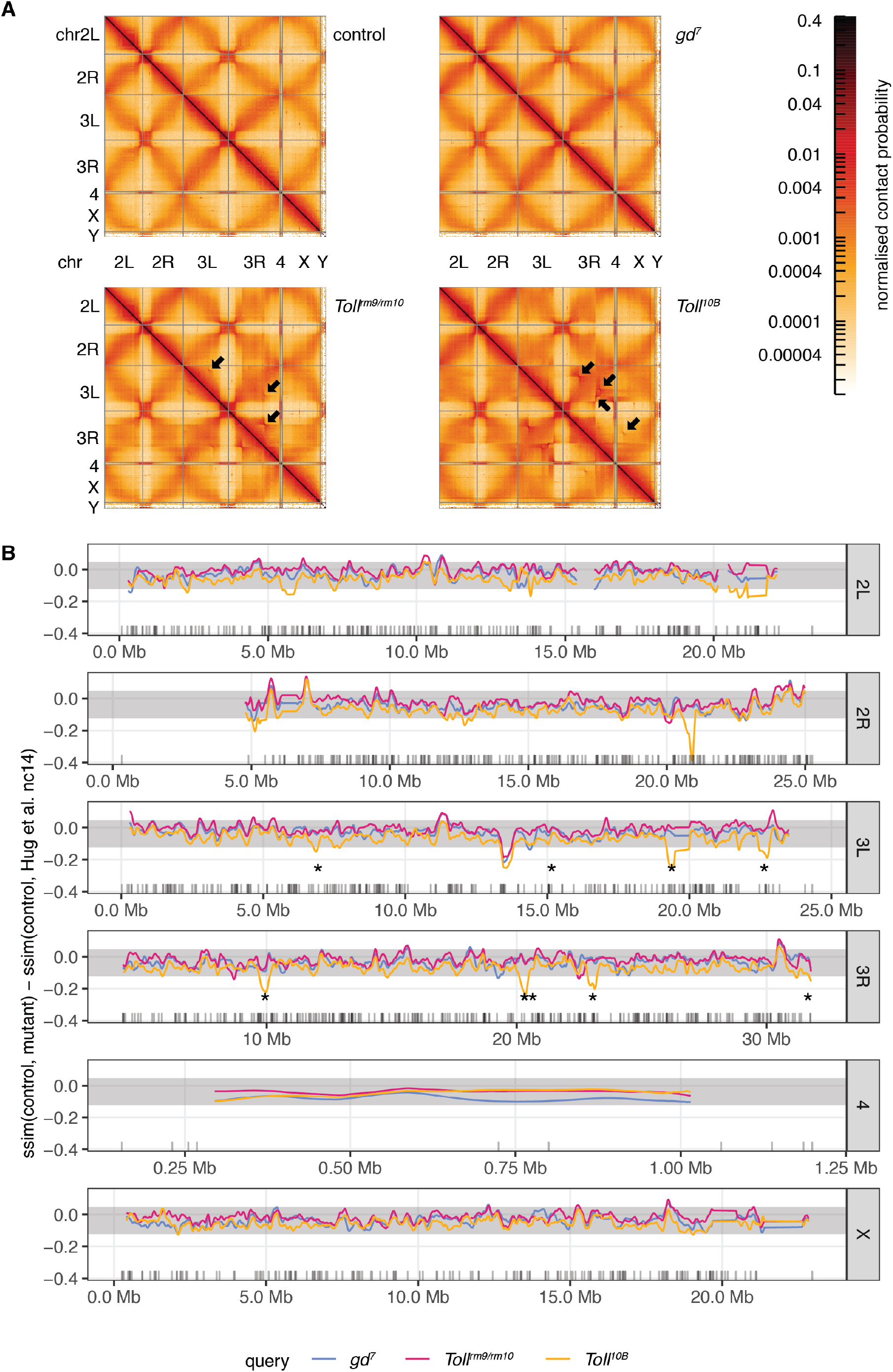
Balancer chromosomes contribute to differences between Hi-C matrices in embryos from different genotypes. **A.** Whole genome contact probability maps for control, *gd^7^*, *Toll^rm9rm/10^*, and *Toll^10B^* embryos at the cellular blastoderm stage. The boundaries of assembled chromosomes and chromosome arms are marked by grey lines. Black arrows mark artefacts in the Hi-C data that indicate rearrangements on balancer chromosomes present in a subset of *Toll^rm9rm/10^* (TM6) and *Toll^10B^* (TM3 and OR60) embryos. **B.** CHESS (Galan, Machnik et al., in revision; see companion manuscript) similarity scores were calculated between mutant and control embryo Hi-C datasets, using 5kb resolution and a 500 kb window size. As a reference, similarity scores were calculated between control embryo Hi-C data and the nc14 Hi-C data from Hug et al. 2017. The difference between this reference similarity score and the similarity scores between mutant and control embryos is shown for all chromosomes (blue, *gd^7^*; pink, *Toll^rm9/rm10^*; yellow, *Toll^10B^*). Similarity score differences around zero represent regions where chromatin conformation is similar between control and mutant, while negative values represent regions where there are greater differences between control and mutant than between control and the Hug et al. nc14 data. Shaded area represents values within two standard deviations of the genome-wide mean. Grey ticks represent the positions of genes that are differentially expressed between dorsoventral mutant embryos (Koenecke et al., 2016). Asterisks mark positions of known rearrangement breakpoints on the TM3 balancer chromosome (Ghavi-Helm et al., 2019).

**Figure S5.**
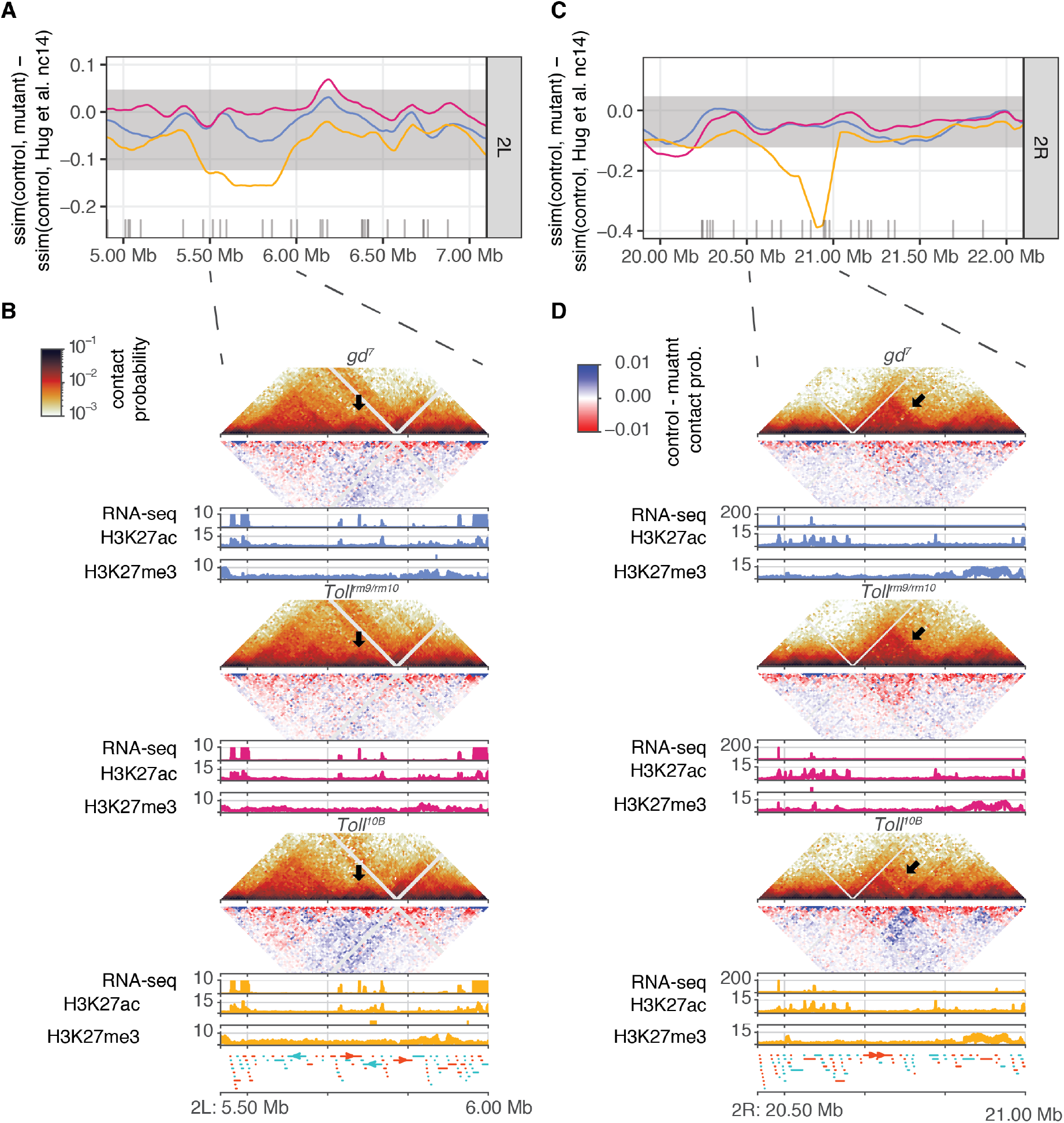
A small number of regions have changes in chromatin conformation detected by CHESS that are not associated with known genomic rearrangements. **A, C.** CHESS (Galan, Machnik et al., in revision; see companion manuscript) similarity scores were calculated between mutant and control embryo Hi-C datasets, using 5kb resolution and a 500 kb window size. As a reference, similarity scores were calculated between control embryo Hi-C data and the nc14 Hi-C data from Hug et al. 2017. The difference between this reference similarity score and the similarity scores between mutant and control embryos is shown for all chromosomes (blue, *gd^7^*; pink, *Toll^rm9/rm10^*; yellow, *Toll^10B^*). Similarity score differences around zero represent regions where chromatin conformation is similar between control and mutant, while negative values represent regions where there are greater differences between control and mutant than between control and the Hug et al. data. Shaded area represents values within two standard deviations of the genome-wide mean. Grey ticks represent the positions of genes that are differentially expressed between dorsoventral mutant embryos (Koenecke et al., 2016). **B, D.** For each genotype, top, normalised Hi-C contact probability maps and Hi-C difference maps at 5kb resolution. Hi-C difference maps are calculated as *contact probability in control – contact probability in mutant*; red indicates regions with increased contact probability in embryos of the mutant genotype and blue indicates decreased contact probability. The black arrows indicate a region with a change in contact probability in *Toll^10B^*. Bottom, RNA-seq (Koenecke et al., 2016), H3K27ac and H3K27me3 ChIP-seq data ((Koenecke et al., 2016), this study). Tissue-specific putative enhancers identified as described above are shown as colour-coded bars beneath the corresponding H3K27ac ChIP-seq track. Lower panel, gene annotations. Positive-strand genes are shown in orange and negative strand genes are shown in blue. **A.** Example of differences in CHESS similarity scores for a 2 Mb region on chromosome 2L. **B.** Hi-C data for a 500 kb subset of the region shown in **A.** **C.** Example of differences in CHESS similarity scores for a 2 Mb region on chromosome 2R. **D.** Hi-C data for a 500 kb subset of the region shown in **C.**

**Figure S6.**
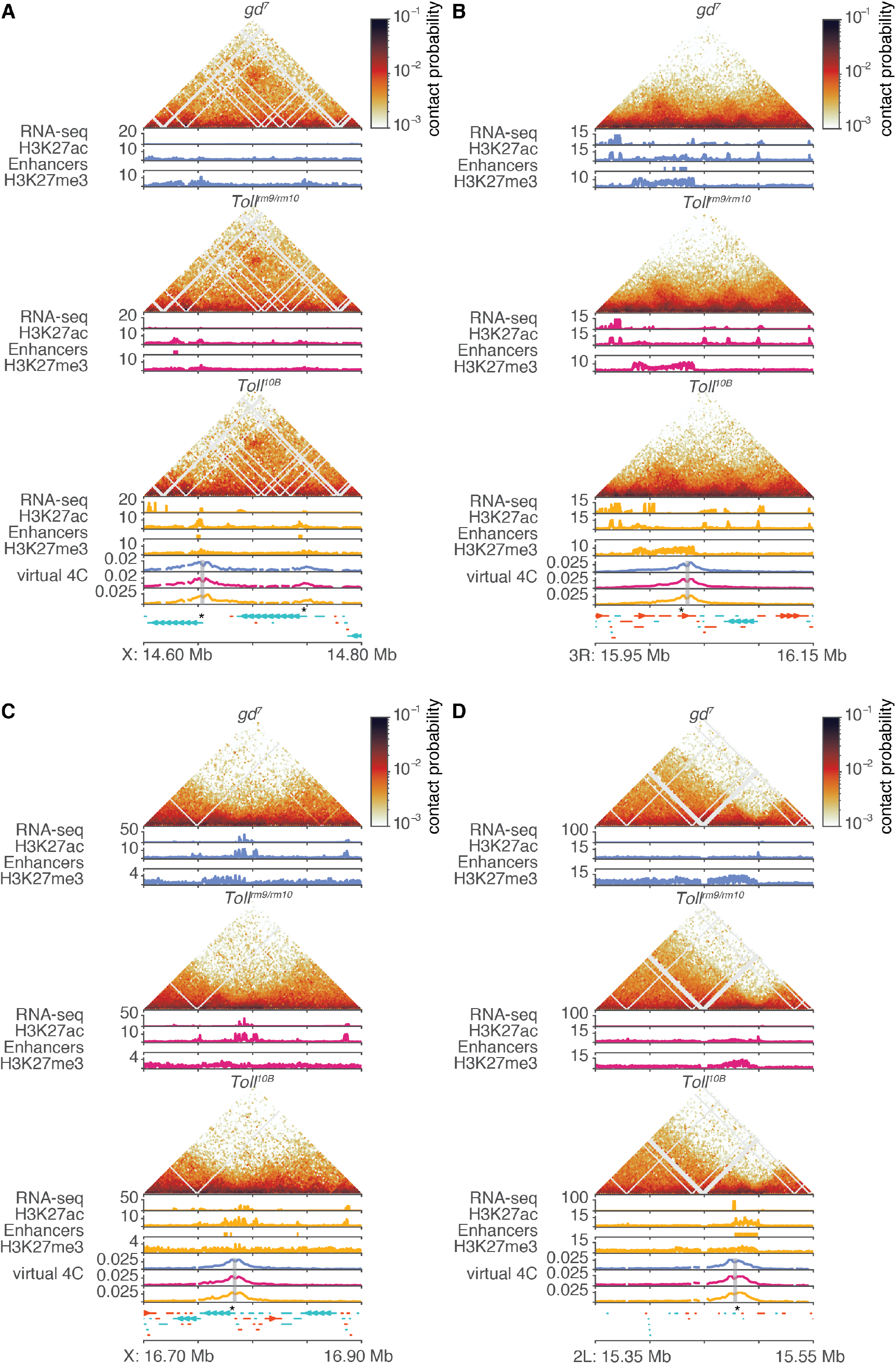
Chromatin conformation is not affected by tissue-specific gene expression. Tissue-specific chromatin data for regions containing dorsoventral patterning genes. For each genotype, top, normalised Hi-C contact probability maps, at 2kb resolution; middle, RNA-seq data (Koenecke et al., 2016); bottom, H3K27ac and H3K27me3 ChIP-seq data ((Koenecke et al., 2016), this study). Tissue-specific putative enhancers identified as described above are shown as colour-coded bars beneath the corresponding H3K27ac ChIP-seq track. Lower panels, “Virtual 4C” tracks for each genotype representing interactions of a 2kb region around the promoters of genes of interest, as highlighted by the grey rectangle; gene annotations. Positive-strand genes are shown in orange and negative strand genes are shown in blue. Dorsoventral patterning genes of interest are marked with asterisks. **A.** *NetA* and *NetB*. **B.** *pnr*. **C.** *if*. **D.** *sna*.

**Figure S7.**
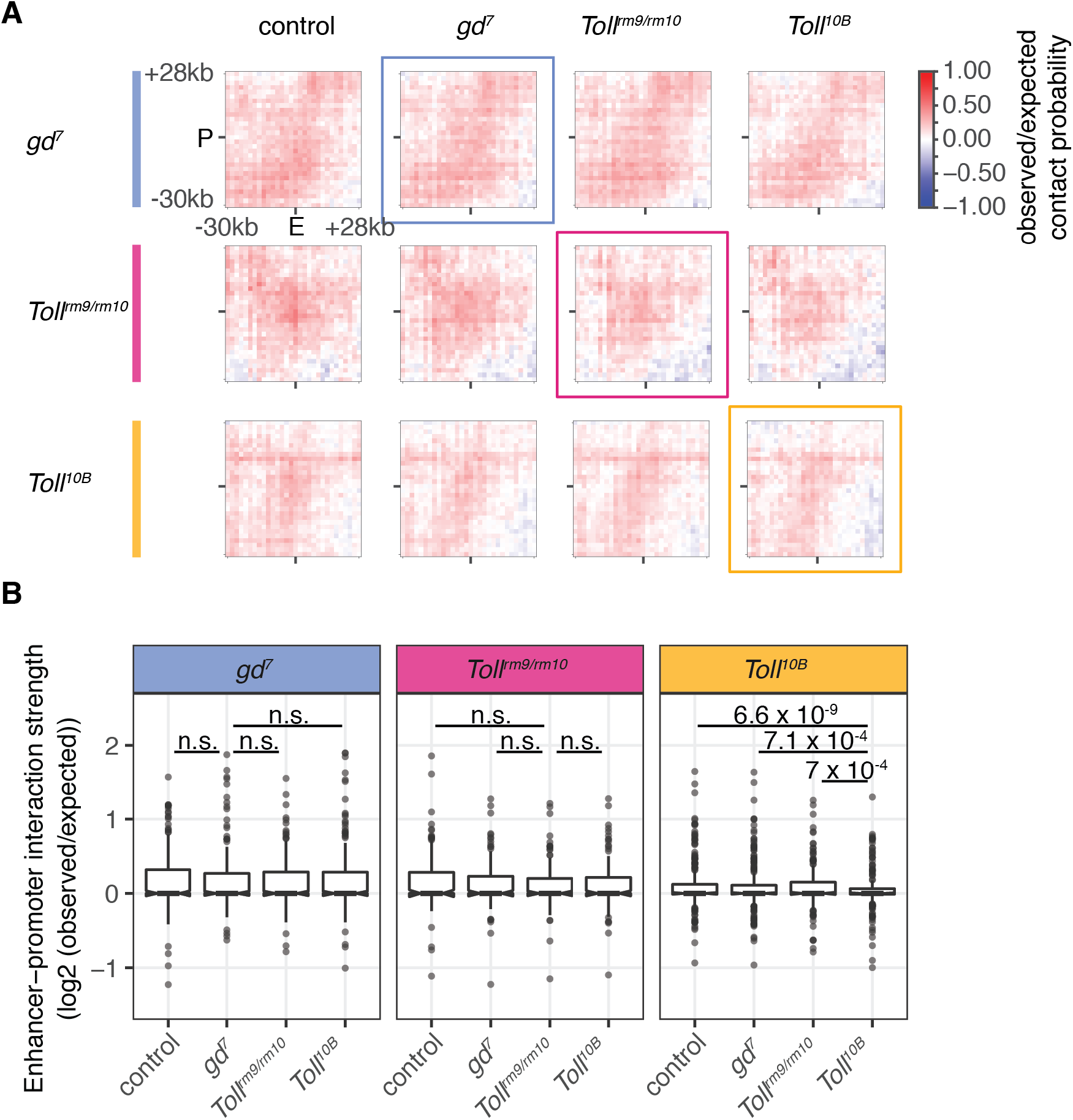
Interaction strength between putative tissue-specific enhancers and the promoters of their assigned genes does not change between tissues. **A.** Aggregate contact analysis for putative tissue-specific enhancers (E) and the promoters (P) of their assigned genes. The average observed/expected contact probability is shown for Hi-C data at 2 kb resolution in a window of 60 kb around putative enhancer-promoter interactions. Rows represent different sets of putative tissue-specific enhancers, while columns represent Hi-C data from cellular blastoderm embryos of different genotypes. Coloured squares highlight datasets in which the enhancers in that row are active (blue, *gd^7^*; pink, *Toll^rm9/rm10^*; yellow, *Toll^10B^*). **B.** Quantification of contact probability between putative enhancers and their assigned target promoters, in Hi-C datasets from embryos of different genotypes. Panels represent different sets of putative tissue-specific enhancers, while the x-axis represents Hi-C data from cellular blastoderm embryos of different genotypes. There are no significant differences in interaction strength across Hi-C datasets for *gd^7^* or *Toll^rm9/rm10^* enhancers. *Toll^10B^* enhancers have significantly lower contacts with their target promoters in Hi-C from *Toll^10B^* mutant embryos.

